# An integrated transcriptomic cell atlas of human neural organoids

**DOI:** 10.1101/2023.10.05.561097

**Authors:** Zhisong He, Leander Dony, Jonas Simon Fleck, Artur Szałata, Katelyn X. Li, Irena Slišković, Hsiu-Chuan Lin, Malgorzata Santel, Alexander Atamian, Giorgia Quadrato, Jieran Sun, Sergiu P. Paşca, J. Gray Camp, Fabian Theis, Barbara Treutlein

## Abstract

Neural tissues generated from human pluripotent stem cells in vitro (known as neural organoids) are becoming useful tools to study human brain development, evolution and disease. The characterization of neural organoids using single-cell genomic methods has revealed a large diversity of neural cell types with molecular signatures similar to those observed in primary human brain tissue. However, it is unclear which domains of the human nervous system are covered by existing protocols. It is also difficult to quantitatively assess variation between protocols and the specific cell states in organoids as compared to primary counterparts. Single-cell transcriptome data from primary tissue and neural organoids derived with guided or un-guided approaches and under diverse conditions combined with large-scale integrative analyses make it now possible to address these challenges. Recent advances in computational methodology enable the generation of integrated atlases across many data sets. Here, we integrated 36 single-cell transcriptomics data sets spanning 26 protocols into one integrated human neural organoid cell atlas (HNOCA) totaling over 1.7 million cells. We harmonize cell type annotations by incorporating reference data sets from the developing human brain. By mapping to the developing human brain reference, we reveal which primary cell states have been generated in vitro, and which are under-represented. We further compare transcriptomic profiles of neuronal populations in organoids to their counterparts in the developing human brain. To support rapid organoid phenotyping and quantitative assessment of new protocols, we provide a programmatic interface to browse the atlas and query new data sets, and showcase the power of the atlas to annotate new query data sets and evaluate new organoid protocols. Taken together, the HNOCA will be useful to assess the fidelity of organoids, characterize perturbed and diseased states and facilitate protocol development in the future.

## Main

Human neural organoids, self-organizing 3D human neural tissues that can be grown in vitro, are becoming powerful tools for studying the mechanisms of human brain development, evolution and disease^1−3^. Neural organoids can be generated using external patterning factors (e.g. morphogens) to guide their development towards certain brain regions or to drive the emergence of specific cell types (guided protocols)^4−8^. Conversely, unguided protocols rely on the self-patterning capacity of organoids to generate diverse cell types and states^9,10^ .

Single-cell RNA sequencing (scRNA-seq) provides a versatile tool to characterize cell type heterogeneity in human neural organoids, and has been used to characterize molecular features of neural cells generated in different neural organoid protocols. It also enables the comparison of human neural organoid cell types and states to those in the primary human brain, and has revealed many similarities in molecular signatures^11^. Nevertheless, there are no comprehensive surveys that catalog all cell states that can arise in neural organoid systems. As a consequence, it is still unclear which portions of the developing central nervous system can already be generated with existing protocols and which ones are still lacking. It has also remained challenging to systematically quantify the transcriptomic fidelity of neural organoid cells compared to their primary counterpart.

In this study, we address these challenges by combining 36 scRNA-seq data sets covering numerous human neural organoid protocols, to generate a transcriptomic cell atlas of human neural organoids using state-of-art computational methods. We establish an analytical pipeline, which allows for the comprehensive and quantitative comparison of the organoid atlas to a recently published reference atlas of the developing human brain^12^. By combining the two atlases, we harmonize annotations of neural and non-neural populations in the two systems, estimate the capacity and precision of different neural organoid protocols to generate neural cells representing different brain regions, and identify primary cell populations that are under-represented in neural organoid protocols. We further estimate transcriptomic fidelity of neurons in neural organoids and identify cell stress as a universal factor distinguishing metabolic states of in vitro neurons from primary neurons. Finally, we map the data of a recent neural organoid morphogen screen^13^ to the integrated atlas to assess regional specificity, and generation of novel states. Together, our work provides a rich atlas resource and a new framework to assess the fidelity of neural organoids, characterize perturbed and diseased states and streamline protocol development in the future.

## Results

### Establishment of a human neural organoid cell atlas (HNOCA) by data curation, harmonization and integration

To build a transcriptomic human neural organoid cell atlas (HNOCA), we collected single-cell RNA sequencing data and detailed, harmonized technical and biological metadata from 36 different data sets, including 33 published^5*−*7,14*−*35^ and three currently unpublished ones (Supplementary Table 1), accounting for 1.77 million cells after consistent preprocessing and quality control (Fig. 1a). The HNOCA represents cell types and cell states that are generated with 26 distinct neural organoid differentiation protocols, including three unguided and 23 guided ones, at different time points ranging from seven to 450 days (Fig. 1b). To remove sample-specific batch effects, we implemented a three-step integration pipeline. First, to anchor our analysis in primary human developing brain data, we projected the HNOCA to a single-cell transcriptomic reference atlas of the developing human brain^12^ using reference similarity spectrum (RSS)^16^. Then, to allow for informed, label-aware integration, we developed an algorithm (snapseed, see Methods) to perform an initial marker-based hierarchical cell type annotation. Last, we used scPoli36 for label-aware data integration based on the hierarchical snapseed annotations. Evaluation of scPoli together with other integration approaches using the previously established benchmarking pipeline^37^ showed that scPoli had the best performance for these data sets (Extended Data Fig. 1). We performed Leiden^38^ clustering based on the scPoli representation and refined annotations for these clusters based on the expression of canonical markers, organoid sample age, as well as the auto-generated cell type labels. A UMAP embedding revealed three neuronal differentiation trajectories corresponding to dorsal telencephalic, ventral telencephalic and non-telencephalic populations as well as trajectories leading from progenitor cells to non-neuronal cell types such as astrocytes and oligodendrocytes precursors (OPCs; Fig. 1c-e; Extended Data Fig. 2). Cells from both unguided and guided protocols were distributed across all trajectories (Fig. 1f).

**Fig. 1.**
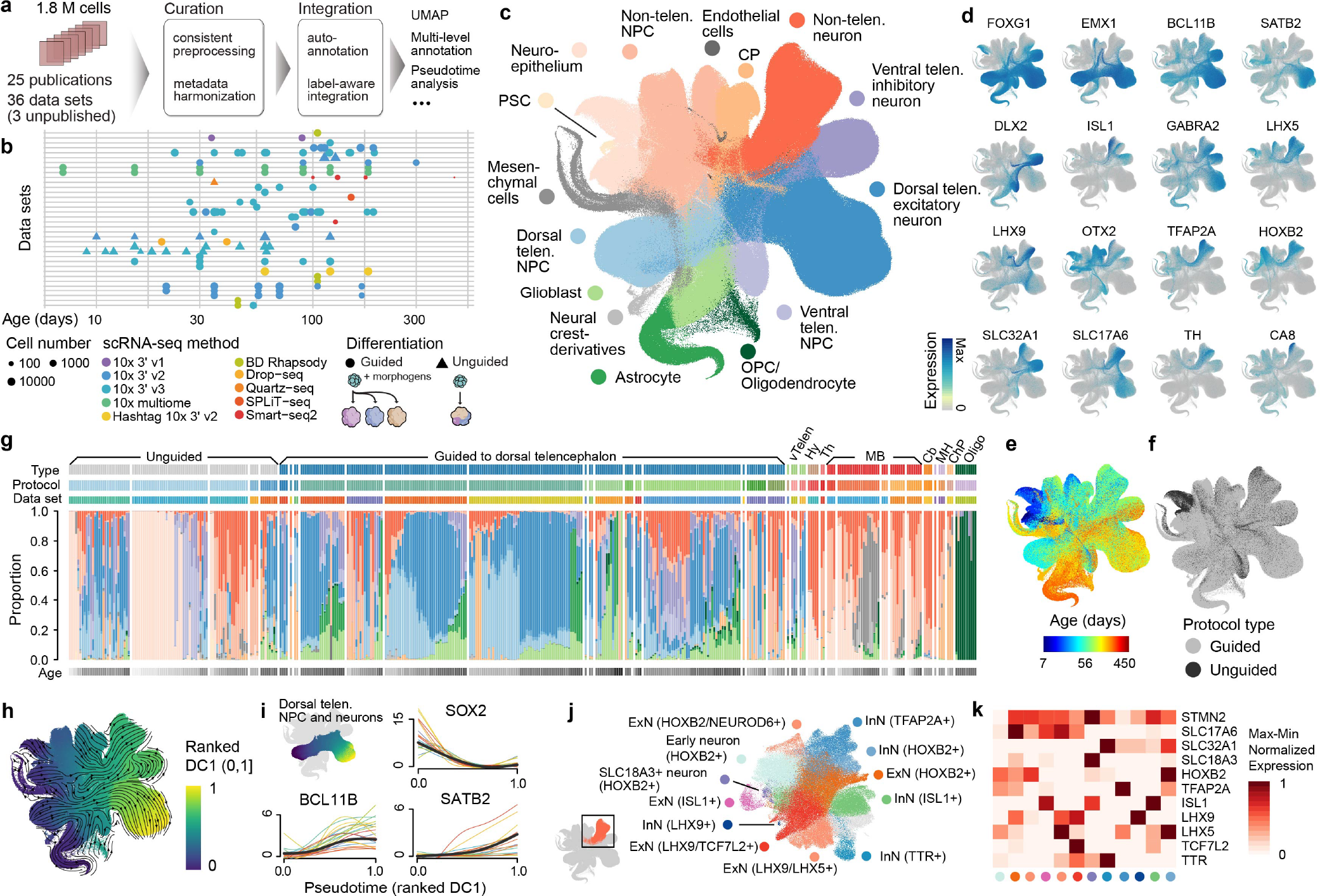
Integrated Human Neural Organoid Cell Atlas (HNOCA). (a) Overview of HNOCA construction pipeline. (b) Metadata of biological samples included in HNOCA. (c-f) UMAP of the integrated HNOCA, colored by (c) level-2 cell type annotations (PSC -pluripotent stem cell, NPC -neural progenitor cell, CP -choroid plexus, OPC -oligodendrocyte progenitor cell, telen. -telencephalon), (d) gene expression profiles of selected regional and cell type markers, (e) sample ages, and (f) organoid differentiation protocol types. (g) Proportions of cells assigned to different cell types (level-2) in HNOCA. Every stacked bar represents one biological sample, grouped into blocks for different data sets and ordered by increasing sample ages. Top bars show the 36 data sets, the used organoid differentiation protocols and protocol types (vTelen -ventral telencephalon, Hy -hypothalamus, Th -thalamus, MB -midbrain, Cb -cerebellum, MH -medulla, ChP -choroid plexus, Oligo -oligodendrocyte). Bottom bars show the sample age. (h) UMAP of the integrated HNOCA colored by top-ranked diffusion component (DC1) on the real-time-informed transition matrix between cells. The stream arrows visualize the inferred flow of cell states toward more mature cells. (i) Expression profiles along the cortical pseudotime of SOX2 (radial glia), BCL11B (deeper layer cortical excitatory neurons) and SATB2 (upper layer cortical excitatory neurons). (j) UMAP of non-telencephalic neurons, colored and labeled by clusters. (k) Heatmap showing relative expression levels of selected genes across different non-telencephalic neuron clusters. The colors of dots at the bottom represent cluster identities as shown in (j).

**Fig. 2.**
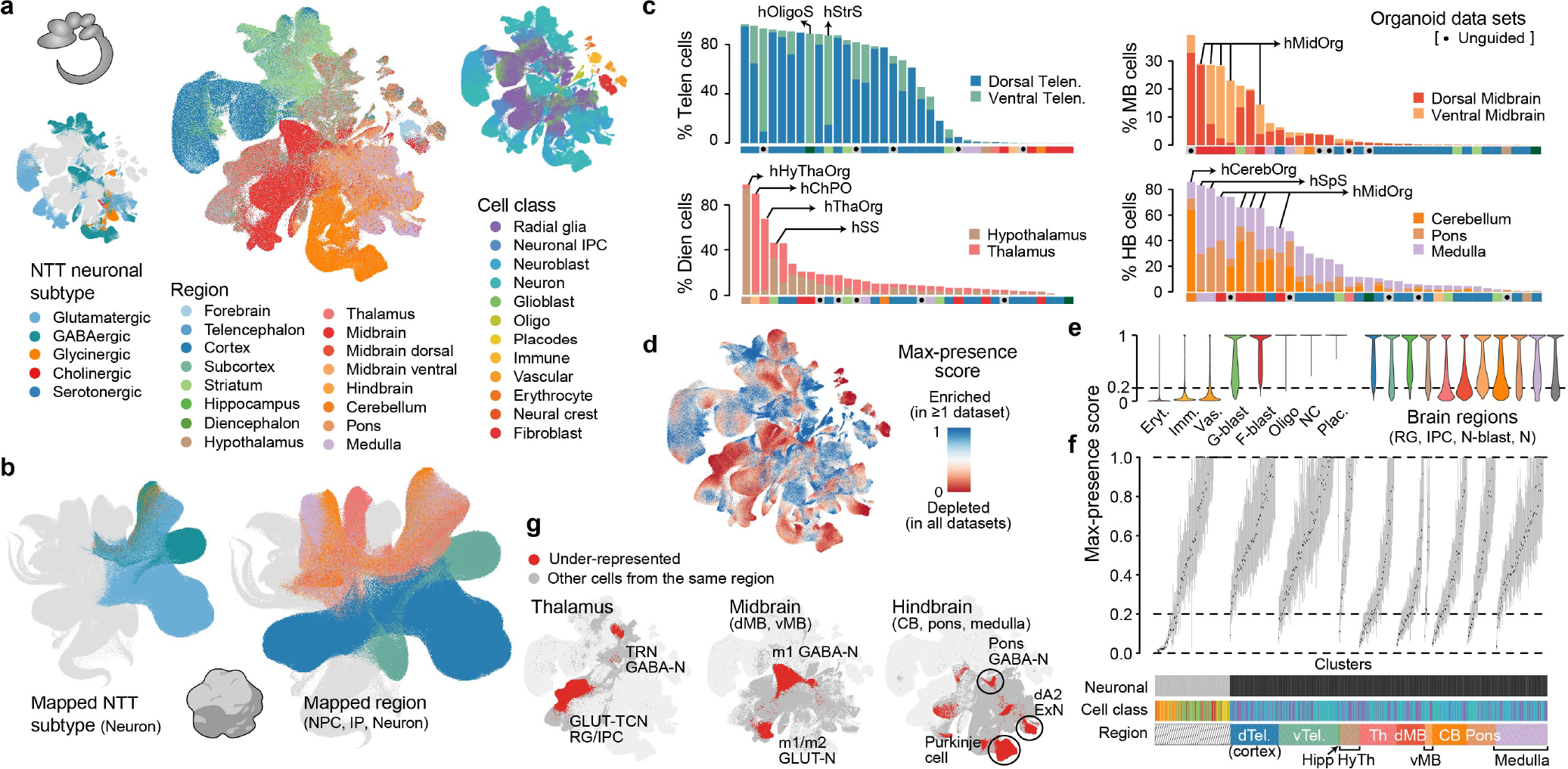
Projection of HNOCA to the primary developing human brain cell atlas assists organoid neural cell type annotation and comprehensive estimation of primary cell type representation in HNOCA. (a) UMAP of the human developing brain cell atlas^12^, colored by neurotransmitter transporter (NTT) subtypes (left), regional labels (middle), and annotated cell classes (right). (b) UMAP of HNOCA, colored by the mapped NTT subtypes of neurons (left) and mapped regional labels of neural progenitor cells (NPCs), intermediate progenitor cells (IP) and neurons. (c) Percentages of neural cells (NPC, IP and neurons) representing different regions, including telencephalon (dorsal and ventral), diencephalon (hypothalamus and thalamus), midbrain (dorsal and ventral), and hindbrain (cerebellum, pons and medulla), in different data sets. The x-axes show data sets, descendingly ordered by the total proportions (bar height). Data sets based on unguided differentiation protocols are marked by dots underneath, with selected guided protocols also labeled. The bars at the bottom of each panel shows organoid protocol type (unguided and guided to different regions). (d) UMAP of the human developing brain cell atlas12 colored by cell population presence within HNOCA data sets (max presence score). A high max presence score suggests enrichment of the corresponding primary cell state in at least one HNOCA data set, with a low score meaning under-representation of the cell state in all HNOCA data sets. (e) Distribution of the max-presence scores of different cell classes in the human developing brain reference atlas12. (f) Box plots showing distribution of max-presence scores in different primary reference cell clusters. The annotation underneath shows cell class and the commonest region information of the primary reference cell clusters. Tel. -telencephalon, Hippo -hippocampus, HyTh -hypothalamus, Th -thalamus, MB -midbrain, d -dorsal, v -ventral. (g) UMAP of the human developing brain atlas showing primary neural cell type/states in thalamus (left), midbrain (middle) and hindbrain (right) that are under-represented in HNOCA (in red).

To elucidate the dynamics and transitions of cell states and cell types represented in the HNOCA, we reconstructed a real-age-informed pseudotime of HNOCA cells based on neural optimal transport (OT) using the moscot framework39 (Fig. 1h). Focusing on dorsal telencephalic neural progenitor cells (NPCs) and neurons, we observed consistent pseudotemporal expression profiles of genes such as NPC marker SOX2, as well as neuronal markers BCL11B (CTIP2) for deeper layer neurons, and SATB2 for upper layer neurons (Fig. 1i). To further resolve heterogeneity among non-telencephalic neurons, we performed sub-clustering of this population revealing numerous neuronal populations characterized by distinct marker gene expression (Fig. 1j-k).

### Projection of the HNOCA to a human primary developing brain reference atlas

To assess our cell type annotation, and more precisely annotate the heterogeneous non-telencephalic neuronal populations, a comprehensive reference atlas of human neural cell types and states is needed. Because neural organoids generally model early stages of the human central nervous system, we used a recently published single-cell transcriptomic atlas of the developing human brain^12^ (Fig. 2a) as the reference for comparison with the HNOCA. We applied scVI^40^ and scANVI^41^ to the primary reference atlas, and used scArches^42^ to project the HNOCA to the same latent space. The shared latent space allowed us to reconstruct a bipartite weighted k-nearest neighbor (wKNN) graph between cells in the HNOCA and the primary reference atlas. With the established wKNN graph, the ‘CellClass’ and ‘Subregion’ labels, as well as the neurotransmitter transporter (NTT) information of neuroblasts and neurons were transferred to the HNOCA. The transferred labels are strongly consistent with our assigned labels (Extended Data Fig. 3) and allowed us to refine the regional annotation of non-telencephalic NPCs and neurons in the HNOCA, as well as the NTT annotation of the non-telencephalic neurons (Fig. 2b), resulting in the final hierarchical cell type annotation of the HNOCA (Extended Data Fig. 3).

**Fig. 3.**
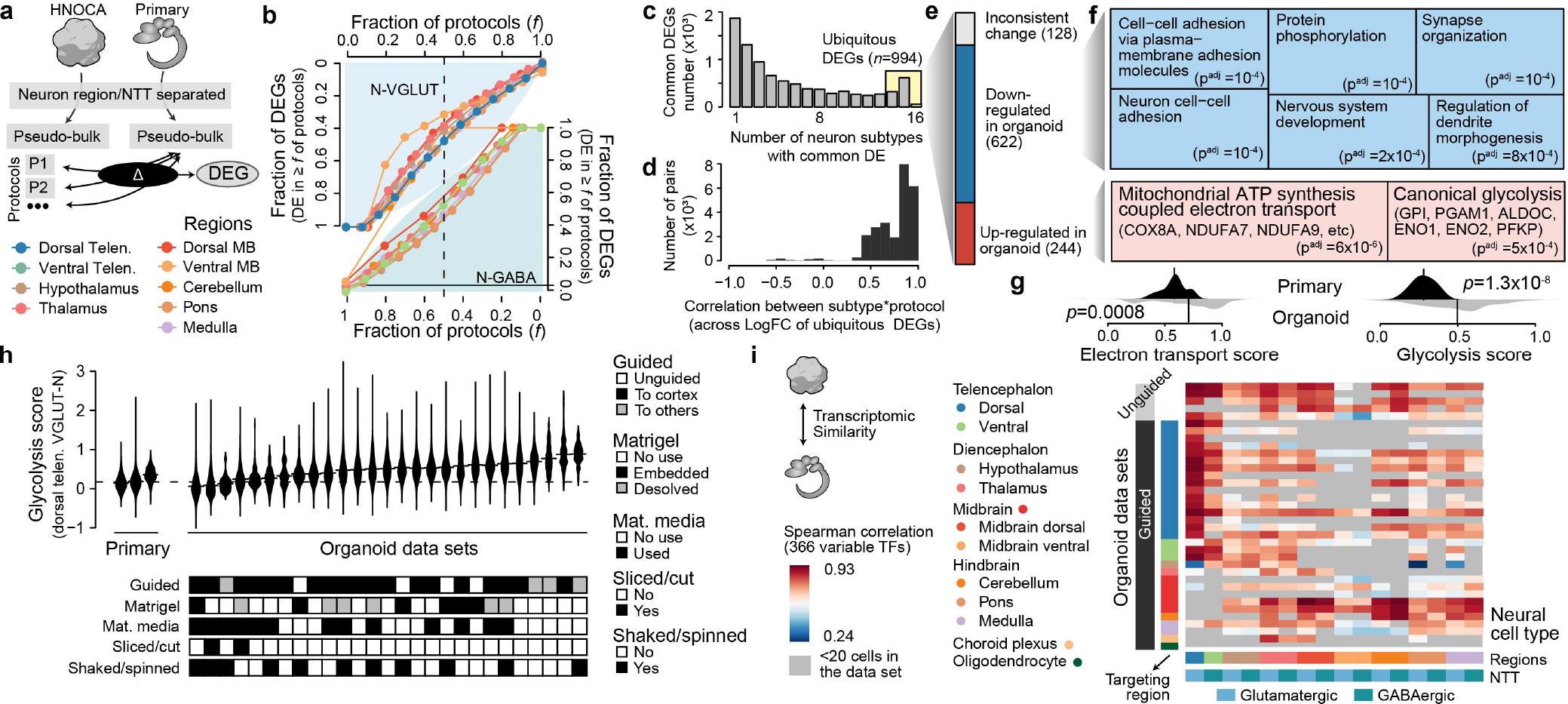
Transcriptomic comparison between organoid neurons and their primary counterpart reveals universal cell stress in organoids. (a) Schematic of differentiation expression (DE) analysis comparing different neural cell types generated by different differentiation protocols in HNOCA to their primary counterpart in the human developing brain atlas^12^. (b) Proportions of expressed genes in different neural cell types that show DE in certain fractions of protocols that generate the corresponding subtypes. The top-left area is for glutamatergic neurons, while the bottom-right area is for GABAergic neurons. Neurons of different regions are colored differently. The x-axes show fractions of protocols, denoted as f. The y-axes show fractions of differentially expressed genes (DEGs) in at least f of all protocols generating the subtypes. The vertical dashed lines denote f=0.5. (c) Numbers of protocol-common DEGs, grouped by the number of neural cell types a gene is estimated to be common DEG. A common DEG in one neural cell type shows DE in at least half of protocols generating the subtype. The 994 genes showing common DE in at least 14 out of 16 neural cell types are defined as ubiquitous DEGs. (d) Distribution of correlation of ubiquitous DEGs log-transformed expression fold change in relative to primary (logFC), between different neural cell types as well as different protocols. (e) Ubiquitous DEGs are classified into three categories: inconsistent change (different subtypes show different DE direction), downregulated, and upregulated in organoids. (f) Gene ontology (GO) enrichment analysis of downregulated (upper, blue) and upregulated (lower, red) ubiquitous DEGs. Sizes of the squares correlate with -log-transformed adjusted p-values. (g) Distribution of gene set (left: mitochondrial ATP synthesis coupled electron transport, electron transport scores; right: canonical glycolysis, glycolysis scores) scores in primary neural cell types (upper, dark), and organoid counterparts (lower, light). P-values show significance of two-sided Wilcoxon test. Both scores are calculated with the score_genes function in scanpy. (h) Glycolysis score, as the proxy of cell stress, of dorsal telencephalic excitatory neurons (dTelen VGLUT-N), split by the three primary developing human brains and 27 organoid data sets with at least 20 dTelen VGLUT-N. Data sets are ordered by their glycolysis score medians. The lower panel shows selected features of differentiation protocols which may be relevant to cell stress. (i) Spearman correlation between gene expression profiles of neural cell types in HNOCA and those in the human developing brain atlas12, across the variable TFs.

Based on the final regional annotation of the HNOCA, we evaluated the capacity of each neural organoid protocol to generate neural cells of different brain regions (Fig. 2c; Extended Data Fig. 3 and 4; Supplementary Table 2). The five data sets of unguided neural organoids show a wide distribution of cells across all brain regions with proportions varying across data sets, indicating the capacity of unguided protocols to generate many brain regions but with high batch-to-batch and line-to-line variability. In contrast, we find that data sets derived from guided organoid protocols are strongly enriched for cells of the targeted brain region. This supports the use of morphogen guidance to efficiently generate a given brain region of interest. Further, we found that data sets of organoids guided to a certain region often show an increased proportion of cells of the brain regions neighboring the targeted regions in the neural axis. For example, several data sets derived from midbrain organoid protocols also show high proportions of diencephalic and hindbrain neurons, indicating an imprecision of morphogen guidance for deep regions of the brain.

**Fig. 4.**
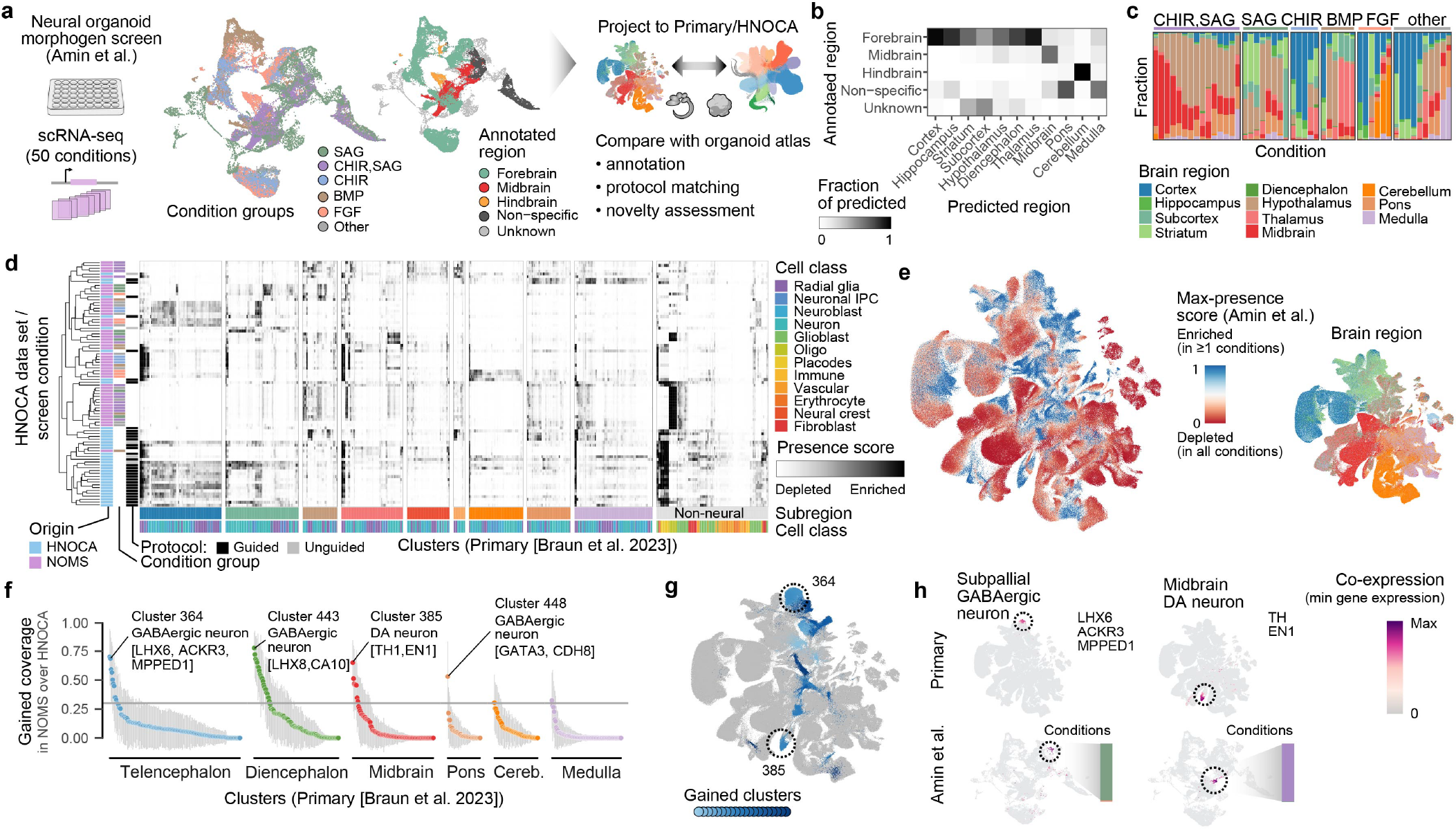
Projection of the neural organoid morphogen screen scRNA-seq data to HNOCA and the human developing brain atlas^12^ allows cell type annotation and organoid protocol evaluation. (a) Schematic of projection of the neural organoid morphogen screen^13^ scRNA-seq data to HNOCA, and the human developing brain atlas^12^ as the primary reference. The UMAPs show screen condition groups (left) and the regional annotation of the screen data (right). (b) Comparison of the regional annotation of the screen data (rows) and the scArches-transferred regional labels from the primary reference. (c) Proportions of cells in the screen data set assigned to different regions by projection to the primary reference. Every stacked bar represents one screened condition. (d) Clustering of HNOCA together with different conditions in the screen data sets, based on the average presence scores of clusters in the primary reference. The dendrogram on the left shows the hierarchical clustering result. The heatmap shows average presence scores per cluster in the primary reference (columns), given data of each protocol involved in HNOCA or the screen condition (rows). (e) UMAP of the primary reference, colored by the dissected regions (left) and the max presence scores across the screen conditions (right). (f) Gain of cell cluster coverage of the screen conditions relative to HNOCA data sets, defined as the difference of average max presence scores per cluster of the primary reference with negative values trimmed to zero, between the screen data set to HNOCA. The gray horizontal line shows the threshold (0.3) to define gained clusters in screen data. (g) UMAP of the primary reference, with gained clusters highlighted in shades of blue. Dashed circles highlight three clusters with the highest gain of coverage in telencephalon, midbrain and hindbrain, respectively. (h) Co-expression scores of cluster marker genes of the three clusters highlighted in (g), in the primary reference (upper) and screen data set (lower).

To evaluate how well organoid protocols represented by the HNOCA generate the different primary cell types or states, we went beyond coarse regional labels and estimated presence scores for every primary cell type in each HNOCA data set (see Methods). A large presence score indicates high frequency and likelihood that cells of a similar type and state are observed in the HNOCA data set. After normalizing the scores per organoid data set (Extended Data Fig. 5; Supplementary Table 3), we obtained the max presence score for each primary reference cell type (Fig. 2d) as the metric to describe how well the cell type is represented in at least one HNOCA data set. This analysis confirmed the absence of erythrocytes, immune cells and vascular endothelial cells in the HNOCA, all of which are derived from non-neuroectoderm germ layers during development (Fig. 2e). The presence of neural cell populations (radial glia (RG), intermediate progenitor cells (IPC), neurons) from different brain regions varied in the organoid data sets. As expected, telencephalic cell types are most strongly represented in HNOCA, as half of the data sets were based on protocols guided towards telencephalon. In contrast, cell types of the thalamus, midbrain and cerebellum are least represented, including thalamic reticular nucleus (TRN) GABAergic neurons, dorsal midbrain m1-derived GABAergic neurons and m1/m2-derived glutamatergic neurons, and cerebellar Purkinje cells (Fig. 2f,g). It is worth noting that, even though these cell types are less abundant in the HNOCA organoid datasets than in the primary atlas, certain organoid protocols are able to generate them (e.g. Purkinje cells generated by cerebellum and midbrain organoid protocols).

### Transcriptomic fidelity of primary neural cell types in neural organoids

We next aimed to understand the transcriptional similarities and differences between organoids generated by distinct differentiation protocols as well as between organoids and primary brain tissue. To this end, we identified differentially expressed genes (DEGs) based on pseudobulk replicates, comparing neural cell types in the HNOCA with their respective primary reference cell type^12^ (Fig. 3a; Supplementary Table 4). Interestingly, we found that for most neural cell types over a third (mean=39.8%, sd=11.5%) of DEGs were shared across at least half of the protocols (protocol-common DEGs), suggesting that a substantial portion of the transcriptomic differences between organoid and primary cells was independent of organoid differentiation protocol (Fig. 3b). To ensure that protocol-common DEGs are not an artifact of using a comprehensive, yet single reference data set of the developing human brain, we verified our results using an additional primary human cortex scRNA-seq data set43 and identified significant overlap with the previously identified protocol-common DEGs (Extended Data Fig. 6; Supplementary Table 5). We further aimed to identify differential transcriptional programs that were not just common across differentiation protocols but also shared across regional neural cell types. We identified a total of 994 ubiquitous DEGs (uDEGs) which were differentially expressed in at least half the differentiation protocols across at least 14 out of the 16 regional neural cell types (Fig. 3c). When quantifying the level of consistency in uDEG fold changes across neuron types and protocols, we observed a correlation higher than 0.8 for over half the pairs, underscoring a highly aligned transcriptomic effect within the uDEGs (Fig. 3d). Out of all 994 uDEGs, 244 genes were consistently upregulated and 622 genes were consistently downregulated, leaving only 128 genes (14%) regulated in inconsistent direction across subtypes or protocols (Fig. 3e).

To identify the biological pathways these three groups of uDEGs are involved in, we ran an enrichment analysis of the Gene Ontology Biological Pathways set^44,45^. For the downregulated uDEGs we identified a clear signature of neurodevelopmental processes including neuron cell-cell adhesion and synapse organization (Fig. 3f). In the up-regulated uDEGs, we observed an enrichment of multiple metabolism-associated terms including mitochondrial ATP synthesis coupled electron transport, a term associated with the oxidative phosphorylation energy pathway, and canonical glycolysis, a metabolic pathway which can also function under anaerobic conditions (Fig. 3f). Using our validation approach across multiple primary cortex reference data sets^12,43^, we were able to confirm these enriched terms. In contrast, only very few to no terms were enriched in the inconsistently differentially regulated genes (Extended Data Fig. 6).

An enrichment of energy-associated pathways has previously been associated with metabolic changes caused by the limitations of the current culture media as well as the supply of nutrients^22,34^. When scoring each of the two predominant upregulated gene sets across the HNOCA and the primary reference atlas12, we found that the canonical glycolysis term showed a more significant separation of organoid and primary cells while also being more unique to the organoid cells (Fig. 3g). Given that metabolic changes are thought to be a cell culture-related phenomenon34 (Extended Data Fig. 7), we decided to use the canonical glycolysis score as the universal proxy for such stress effects in organoid cells.

Next, we wanted to explore the cell type specificity of increased glycolysis in organoids. Using the Kanton et al.^16^ and Braun et al.^12^ data sets as representative examples, we identified a similar distribution of glycolysis scores across all neural cell types with an overall increased score in organoid cells (Extended Data Fig. 7). Focusing on dorsal telencephalic neurons, which were present in organoids from a particularly large number of differentiation protocols, we compared the distribution of glycolysis scores across differentiation protocols and identified several protocol features that correlated with metabolic cell stress. For instance, the usage of maturation media, slicing/cutting of organoids and, to a lesser extent, shaking/spinning of organoids led to overall lower glycolysis scores (Fig. 3h).

When comparing mean glycolysis scores and transcriptomic similarities of organoid and primary reference cell types^12^ across differentiation protocols, we observed an overall negative correlation^22,34^. Interestingly, the correlation strength was significantly reduced when considering only variable transcription factors (TFs) in the transcriptomic similarity analysis, indicating that the metabolic changes in organoids has only limited impact on the core molecular identity of neuronal cell types as defined by TF activity (Extended Data Fig. 7).

In order to quantify the transcriptomic similarity between organoid and primary reference cell types^12^ while avoiding the confounding effect of metabolic changes in organoids, we focused on the expression of 366 variable TFs to calculate the correlation between corresponding neuronal cell types in the HNOCA data sets and the primary reference atlas12. We found that both guided and unguided organoid differentiation protocols generated neuronal cell types with comparable similarity to the corresponding primary reference cell types. However, we observed brain region dependent differences in transcriptomic similarity between organoid and primary reference neuron types. For example, organoid neurons from the dorsal part of most brain regions such as the dorsal telencephalon, dorsal midbrain and cerebellum showed higher similarity to their primary counterparts across organoid data sets than cell types derived from the ventral part of most brain regions such as the hypothalamus, ventral midbrain and pons (Fig. 3i).

### The HNOCA facilitates evaluation of new neural organoid protocols

The integrated HNOCA, as well as the analytical pipeline we established to compare to the primary reference atlas in terms of cell type composition and transcriptomic fidelity, provide a framework to query novel neural organoid scRNA-seq data sets not included in the HNOCA. To showcase this application, we retrieved the scRNA-seq data of a recently published multiplexed neural organoid morphogen screen^13^, and projected it to the same latent spaces as the HNOCA and the primary reference^12^, respectively, using scArches (Fig. 4a, Extended Data Fig. 8; Supplementary Table 6). Regional labels were predicted for query cells by label transfer from the primary reference using the same procedure as mentioned above. The predicted regional labels were highly consistent with the provided regional annotation, but with higher resolution within each of the broad brain sections of fore-brain, midbrain and hindbrain (Fig. 4b). Our transferred annotation therefore allowed a more comprehensive assessment of the effects of different morphogen conditions on generating neurons of different brain regions (Fig. 4c). We further calculated presence scores for each primary reference cell in each of the screen conditions and compared the data of the different screen conditions with the 36 HNOCA data sets. We used hierarchical clustering on average presence scores of primary reference atlas clusters for this comparison (Fig. 4d). Many screen conditions showed distinct presence score patterns, suggesting the generation of organoids with regional cell type compositions that are distinct from the HNOCA data sets. Next, we summarized the max presence scores for the whole morphogen screen data set (Fig. 4e), and compared then to the max presence scores calculated for the HNOCA data to assess which primary reference cell types showed an increased presence in the screening data (Fig. 4f). This analysis highlighted several reference cell clusters that showed a significant abundance increase under certain screen conditions (Fig. 4g). The top hits include the LHX6/ACKR3/MPPED1 triple-positive GABAergic neurons in the ventral telencephalon, the dopaminergic neurons in ventral midbrain, and Purkinje cells in the cerebellum. In summary, the projection of the morphogen screen query data to the HNOCA and the primary reference atlas allowed a refined annotation of the morphogen screen data, as well as a comprehensive and quantitative evaluation of the value of novel neural organoid protocols to generate neuronal cell types previously underrepresented or lacking in neural organoids.

## Discussion

In this study, we built the first large-scale integrated cell atlas of human neural organoids, the HNOCA. To effectively integrate the 1.8 million cells spanning 36 scRNA-seq data sets generated by 15 different laboratories worldwide using 26 different differentiation protocols as well as diverse scRNA-seq technologies, we introduced a multi-step data integration procedure: 1) pre-integration using RSS^16^ ; 2) marker-based auto-annotation using snapseed; and 3) label-aware integration using scPoli^36^. The resulting integrated atlas revealed the high complexity of neuronal, glial and non-neural cell types that can develop in neural organoids grown under existing protocol conditions. Mapping the HNOCA data to a recently published human developing brain cell reference atlas^12^ allowed comprehensive evaluation of neural organoid protocols to generate cell types of different brain regions, and revealed primary brain tissue cell types that are under-represented in organoids from currently available protocols. We performed differential expression analysis between organoid neuron types and their primary counterparts to evaluate the transcriptomic fidelity, and identified metabolic stress as a main factor that distinguishes organoid and primary cell states. Last, we showcased the mapping of a query data set, a recently published single-cell transcriptomic neural organoid morphogen screen, to the HNOCA and the primary reference, which enabled a refined cell type annotation, as well as a compositional comparison with existing neural organoid data sets. The integrated HNOCA, as well as the entire analytical pipeline, are publicly available, and the variational autoencoder (VAE) used for data integration flexibly enables incorporation of new data sets for future atlas expansion. This powerful framework will facilitate quantitative and comparative analysis of scRNA-seq data of human neural organoids, and for the benchmarking of novel neural organoid protocols.

Consistent with earlier reports^16,46^, we find that unguided protocols generate neural cells with high brain regional variability, which is useful when studying broader fate determination during nervous system development. Guided protocols resulted in a strong enrichment of cell types of the targeted brain regions. We also note that guided protocols, particularly those targeting non-telencephalic regions such as the midbrain, show relatively low specificity and generate neural cells from the nearby brain regions. This issue of broader specification may be due to a differential response of neural stem cells in the organoid to the same morphogen cue, or to the lack of a full understanding of the timing, concentration and combinations of morphogens that would be required to precisely define cells of the deeper regions in the central nervous system. Further work is needed to fully understand which factors control the emergence of any given brain region.

Through comparison to primary developing human brain tissue, we found that metabolic changes related to the glycolysis pathway in in vitro versus primary tissue. Despite the negative effects of metabolic stress on overall transcriptomic fidelity, the molecular identity of regional cell types is maintained as evidenced by transcription factor co-expression patterns that are highly consistent with primary counterparts. We find that transcriptomic fidelity is higher for some neural cell types (e.g. telencephalic neurons) than others (e.g. cerebellar neurons). This could be due to the more distinct molecular signatures of telencephalic neurons relative to others, or the rostral telencephalic neurons being the default fate of early embryonic ectodermal cells as previously proposed^47^ .

Altogether, an integrated organoid atlas will enable us to assess the fidelity of organoids, characterize perturbed and diseased states and streamline protocol development in the future.

## Methods

### Metadata curation and harmonization of human neural organoid scRNA-seq data sets

We included 33 human neural organoid data from a total of 25 publications^5*−*7,14*−*35^ plus three unpublished data sets in our atlas (Supplementary Table 1). We curated all neural organoid data sets used in this study via the sfaira^48^ framework (GitHub dev branch, 18th April 2023). For this, we obtained scRNA-seq count matrices and associated metadata from the location provided in the data availability section for every included publication or directly from the authors in case of unpublished data. We harmonized metadata according to the sfaira standards (https://sfaira.readthedocs.io/en/latest/adding_datasets.html) and manually curated an additional metadata column “organoid_age_days” which described the number of days the organoid had been in culture before collection.

We next removed any non-applicable subsets of the published data sets: diseased samples/samples expressing disease-associated mutations (Huang et al.^7^, Sawada et al.^25^, Khan et al.^26^, Bowles et al.^27^, Samarasinghe et al.^29^, Paulsen et al.^35^), fused organoids (Birey et al.^14^), primary fetal data (Bhaduri et al.^22^, Uzquiano et al.^33^), hormone-treated samples (Kelava et al.^32^), data collected prior to neural induction (Kanton et al.^16^, Fleck et al.^30^), and share-seq data (Uzquiano et al.^33^). We harmonized all remaining data sets to a common feature space using any genes of the biotype “protein_coding” or “lncRNA” from ensembl^49^ release 104 while filling any genes missing in a data set with zero counts. We then concatenated all remaining data sets to create a single AnnData^50^ object.

### Preprocessing of the human neural organoid cell atlas (HNOCA) scRNA-seq data

All processing and analysis was carried out using scanpy^51^ (v1.9.3) unless indicated otherwise. For quality control and filtering of HNOCA, we removed any cells with less than 200 genes expressed. We next removed outlier cells in terms of two quality control (QC) metrics: the number of expressed genes and percentage mitochondrial counts. To define outlier cells based on each QC metric, z-transformation is firstly applied to values across all cells. Cells with any z-transformed metric < -1.96 or > 1.96 are defined as outliers. For any data set collected using the v3 chemistry by 10x Genomics, which contains more than 500 cells after the filtering, we fitted a Gaussian distribution to the histogram denoting the number of expressed genes per cell. If a bimodal distribution was detected, we removed any cell with fewer genes expressed than defined by the valley between the two maxima of the distribution. We then normalized the raw read counts for all Smart-seq2 data by dividing it by the maximum gene length for each gene obtained from BioMart. We next multiplied these normalized read counts by the median gene length across all genes in the data sets and treated those length-normalized counts equivalently to raw counts from the data sets obtained with the help of unique molecular identifiers (UMIs) in our downstream analyses.

As a next step we generated a log-normalized expression matrix by first dividing the counts for each cell by the total counts in that cell and multiplying by a factor of 1,000,000 before taking the natural logarithm of each count+1. We computed 3000 highly variable features in a batch-aware manner using the scanpy highly_variable_genes function (flavor=“seurat_v3”, batch_key=“bio_sample”). Here, “bio_sample” represents biological samples as provided in the original metadata of the data sets. We used these 3000 features to compute a 50-dimensional representation of the data using Principal Component Analysis (PCA) which in turn we used to compute a k-nearest-neighbor (kNN) graph(n_neighbors=30, metric=“cosine”). Using the neighbor graph we computed a two dimensional representation of the data using Uniform Manifold Approximation and Projection (UMAP)^52^ and a coarse (resolution=1) and fine (resolution=80) clustering of the unintegrated data using Leiden38 clustering.

### Hierarchical auto-annotation with snapseed

To obtain initial annotations for label-aware integration we devised a scalable auto-annotation strategy. First, we constructed a hierarchy of cell types including progenitor, neuron and non-neural types, each defined by a set of marker genes (Supplementary Data 1). Next, we represented the data by the reference similarity spectrum (RSS)^16^ to average expression profiles of cell clusters in the recently published human developing brain cell atlas^12^. We then constructed a kNN graph (k=30) in the RSS space and clustered the data set using the Leiden algorithm38 (resolution=80). For both steps we used the GPU-accelerated RAPIDS implementation which is provided through scanpy^51,53^ .

For all cell type marker genes on a given level in the hierarchy, we computed the area under the receiver operating characteristic curve (AUROC) as well as the detection rate across clusters. For each cell type, a score was computed by multiplying the maximum AUROC with the maximum detection rate among its marker genes. Each cluster was then assigned to the cell type with the highest score. This procedure was performed recursively for all levels of the hierarchy. The same procedure was carried out using the fine (resolution=80) clustering of the unintegrated data to obtain cell type labels for the unintegrated data set that were used downstream as a ground-truth input for benchmarking integration methods.

This auto-annotation strategy was implemented in the snapseed python package and is available on GitHub (https://github.com/devsystemslab/snapseed). Snapseed is a light-weight package to enable scalable marker-based annotation for atlas-level data sets, where manual annotation is not readily feasible. The package implements three main functions: annotate() for non-hierarchical annotation of a list of cell types with defined marker genes, annotate_hierarchy() for annotating more complex, manually defined cell type hierarchies and find_markers() for fast discovery of cluster-specific features. All functions are based on a GPU-accelerated implementation of AUROC scores using JAX (https://github.com/google/jax).

### Label-aware data integration with scPoli

We performed integration of the organoid data sets for HNOCA using the scPoli^36^ model from the scArches^54^ package. We defined the batch covariate for integration as a concatenation of the data set identifier (annotation column “id”), the annotation of biological replicates (annotation column “bio_sample”) as well as technical replicates (annotation column “tech_sample”). This resulted in 396 individual batches. The batch covariate is represented in the model as a learned vector of size 5. We used the top three levels of the RSS-based snapseed cell type annotation as the cell type label input for the scPoli prototype loss. We chose the hidden layer size of the one-layer scPoli encoder and decoder as 1024, and the latent embedding dimension as 10. We used a value of 100 for the “alpha_epoch_anneal” parameter. We did not use the unlabeled prototype pretraining. We trained the model for a total of 7 epochs, 5 of which were pre-training epochs.

### Benchmark of data integration methods

To quantitatively compare the organoid atlas integration results from multiple tools, we used the GPU-accelerated scib-metrics^37,55^ python package (v0.3.3) and used the embedding with the highest overall performance for all downstream analyses. We compared the data integration performance across the following latent representations of the data: unintegrated PCA, RSS^16^ integration, scVI^40^ (default parameters except for using 2 layers, latent space of size 30 and negative binomial likelihood) integration, scANVI^41^ (default parameters) integrations using either snapseed level 1, 2 or 3 annotation as cell type label input, scPoli^36^ (parameters shown above) integrations using either snapseed level 1, 2 or 3 annotation or all three annotation levels at once as cell type label input, scPoli36 integrations of meta-cells aggregated with the aggrecell algorithm (first employed as “pseudocell”^16^) using either snapseed level 1 or 3 annotation as cell type label input to scPoli. We used the following scores for determining integration quality (each described in Luecken et al.^37^): Leiden normalized mutual information score, Leiden adjusted rand index, average silhouette width (ASW) per cell type label, Isolated label score (ASW-scored) and cell-type local inverse Simpson’s index to quantify conservation of biological variability. To quantify batch effect removal, we used average silhouette width per batch label, integration local inverse Simpson’s index, k-nearest-neighbor batch-effect test (kBET) score and graph connectivity. Integration approaches were then ranked by an aggregate total score of individually normalized (into the range of [0,1]) metrics. Before we carried out the benchmarking, we iteratively removed any cells from the data set that had an identical latent representation as another cell in the data set until no latent representation contained any more duplicate rows. This procedure removed a total of 3293 duplicate cells (0.002 % of the whole data set) and was required for the benchmarking algorithm to complete without errors. We used the snapseed level 3 annotation computed on the unintegrated PCA embedding as ground truth cell type labels in the integration.

### Pseudotime inference

To infer a global ordering of differentiation state, we sought to infer a real time-informed pseudotime based on neural optimal transport in the scPoli latent space. We first grouped organoid age in days into seven bins ((0, 15], (15, 30], (30, 60], (60, 90], (90, 120], (120, 150], (150, 450]). Next we used moscot^39^ to solve a temporal neural problem. To score the marginal distributions based on expected proliferation rates, we obtained proliferation and apoptosis scores for each cell with the method score_genes_for_marginals(). Marginal weights were then computed with

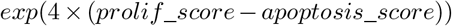

The optimal transport problem was solved using the following parameters: iterations=25000, compute_wasserstein_baseline=False, batch_size=1024, patience=100, pretrain=True, train_size=1. To compute displacement vectors for each cell in age bin *i*, we used the subproblem corresponding to the [*i*, − *i* + 1] transport map, except for the last age bin, where we used the subproblem [*i* − 1, *i*]. Displacement vectors were obtained by subtracting the original cell distribution from the transported distribution. Using the velocity kernel from CellRank^56^ we computed a transition matrix from displacement vectors and used it as an input for computing diffusion maps^57^. Ranks on negative diffusion component 1 were used as a pseudotemporal ordering.

### Preprocessing of the human developing brain cell atlas scRNA-seq data

The cellranger-processed scRNA-seq data for the primary atlas^12^ was obtained from the link provided on its GitHub page (https://storage.googleapis.com/linnarsson-lab-human/human_dev_GRCh38-3.0.0.h5ad). For additional quality control, cells with fewer than 300 detected genes were filtered out. Transcript counts were normalized by the total number of counts for that cell, multiplied by a scaling factor of 10,000 and subsequently natural-log transformed. The feature set was intersected with all genes detected in the organoid atlas and the 2000 most highly variable genes were selected with the scanpy function *highly_variable_genes* using ‘Donor’ as the batch key. An additional column of ‘neuron_ntt_label’ was created to represent the automatic classified neural transmitter transporter subtype labels derived from the ‘AutoAnnotation’ column of the cell cluster metadata (https://github.com/linnarsson-lab/developing-human-brain/files/9755350/table_S2.xlsx).

### Reference mapping of the organoid atlas to the primary atlas

To compare our organoid atlas with data from the primary developing human brain, we used scArches^54^ to project it to the above mentioned primary human brain scRNA-seq atlas^12^. We first pretrained a scVI model^40^ on the primary atlas with ‘Donor’ as the batch key. The model was constructed with following parameters: n_latent=20, n_layers=2, n_hidden=256, use_layer_norm=“both”, use_batch_norm=“none”, encode_covariates=True, dropout_rate=0.2 and trained with a batch size of 1024 for a maximum or 500 epochs with early stopping criterion. Next the model was fine-tuned with scANVI^41^ using ‘Subregion’ and ‘CellClass’ as cell type labels with a batch size of 1024 for a maximum of 100 epochs with early stopping criterion and n_samples_per_label=100. To project the organoids atlas to the primary atlas, we used the scArches^54^ implementation provided by scvi-tools^58,59^. The query model was finetuned with a batch size of 1024 for a maximum of 100 epochs with early stopping criterion and weight_decay=0.0.

### Bipartite weighted k-nearest neighbor (w-kNN) graph reconstruction

With the primary reference^12^ and query (HNOCA) data projected to the same latent space, an unweighted bipartite k-nearest neighbor (kNN) graph was constructed, by identifying 100 NNs of each query cell in the reference data with either PyNNDescent or RAPIDS-cuML (https://github.com/rapidsai/cuml) in Python, depending on availability of GPU acceleration. Similarly, a reference kNN graph was also built by identifying 100 NNs of each reference cell in the reference data. For each edge in the reference-query bipartite graph, the similarity between the reference neighbors of the two linked cells is represented by the Jaccard index:

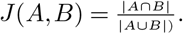

The square of Jaccard index was then assigned as the weight of the edge, to get the bipartite weighted kNN graph between reference and query data sets.

### W-KNN-based primary developing brain atlas label transfer to HNOCA cells

Given the w-KNN estimated between primary reference^12^ and query (HNOCA), any categorical metadata label of reference can be transferred to query cells via majority voting. In brief, for each category, its support was calculated for each query cell as the sum of weights of edges, which link to reference cells in this category. The category with the largest support was assigned to the query cell.

To get the final regional labels for the non-telencephalic neural progenitor cells (NPCs) and neurons, as well as the neurotransmitter transporter (NTT) labels for non-telencephalic neurons, constraints were added to the transfer procedure. For regional labels, only the non-telencephalic regions with the finest resolution, namely hypothalamus, thalamus, midbrain dorsal, midbrain ventral, cerebellum, pons and medulla, were considered as valid categories to be transferred. For NTT labels, we first identified valid region-NTT label pairs in the reference based on the provided NTT labels in the reference neuroblast and neuron clusters and their most common regions. Here, the most common regions were re-estimated in a hierarchical manner, to the finest resolution mentioned above. Next, when transferring NTT labels, for each non-telencephalic neuron with the regional label transferred, only NTT labels that are considered valid for the region were considered during majority voting.

### Presence scores and max presence scores of cells in the primary developing brain atlas

Given a reference data set and a query data set, the presence score is a score assigned to each cell in the reference, which describes the frequency or likelihood of the cell type/state of that reference cell appearing in the query data. In this study, we calculated the presence scores of primary atlas cells in each HNOCA data set to quantify how frequently we saw a cell type/state represented by each primary cell in each of the HNOCA data sets.

Specifically, for each HNOCA data set, we first subset the w-KNN graph to only HNOCA cells in that data set. Next, the raw weighted degree was calculated for each cell in the primary atlas, as the sum of weights of the remaining edges linked to the cell. A random walk with restart (RWWR) procedure was then applied to smooth the raw scores across the kNN graph of the primary atlas. In brief, we first represented the primary atlas kNN graph as its adjacency matrix (*A*), followed by row normalization to convert it into a transition probability matrix (*P*). With the raw scores represented as a vector *s*_0_, in each iteration *t*, we generated *s*_*t*_ as

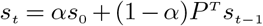

This procedure was performed 100 times to get the smooth presence scores which were subsequently log transformed. Scores lower than the 5^*th*^ percentile or higher than the 95^*th*^ percentile were trimmed. The trimmed scores were normalized into the range of [0, 1] as the final presence scores in the HNOCA data set.

Given the final presence scores in each of the HNOCA data sets, the max presence scores in the whole HNOCA data was then easily calculated as the maximum of all the presence scores for each cell in the primary atlas. A large (close to one) max presence score indicates a high frequency of appearance for the cell type/state in at least one HNOCA data set while a small (close to zero) max presence score suggests under-representation in all the HNOCA data sets.

### Cell type composition comparison among morphogen usage using scCODA

To test the cell type compositional changes upon admission of certain morphogens from different organoid differentiation protocols, we used the pertpy^60^ implementation of the scCODA algorithm^61^. scCODA is a Bayesian model to detect compositional changes in scRNA-seq data. For this, we have extracted the information about the added morphogens from each differentiation protocol and grouped them into 15 broad molecule groups based on their role in neural differentiation (Supplementary Table S1). These molecule groups were used as a covariate in the model. The region labels transferred from the primary atlas were used as labels in the analysis (cell_type_identifier). For cell types without regional identity, the coarse level 1 cell type labels were used. Pluripotent stem cells (PSC) and neuroepithelium cells were removed from the analysis since they are mainly present in the early organoid stages. We used bio_sample as the sample_identifier. We ran scCODA sequentially with default parameters, using No-U-turn sampling (run_nuts function) and selecting each cell type once as a reference. We used a majority vote-based system to find the cell types that were credibly changing in more than half of the iterations.

### Cell type composition comparison among morphogen usage using regularized linear regression

To complement the composition analysis conducted with scCODA, we devised an alternative approach to test for differential composition using regularized linear regression. We fit a generalized linear model with the region composition matrix as the response *Y* and molecule usage as independent variables *X*:

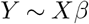

The model was fit with lasso regularization (alpha = 1) using Gaussian noise and an identity link function. The regularization parameter lambda was automatically determined through cross-validation as implemented in the function cv.glmnet() from the glmnet^62^ R package. All non-zero coefficients *β* were considered as indications of enrichment and depletion.

### Differential expression analysis between HNOCA neural cell types and their primary counterparts and functional enrichment analysis

To study the transcriptomic differences between organoid and primary cells, we subset HNOCA using the final level 1 annotation to cells labeled “Neuron”. We furthermore subset the human developing brain atlas to cells that had been assigned a valid label in the “neuron_ntt_label” annotation column. We added an additional two data sets of fetal cortical cells from Camp et al.^11^ and from Eze et al.^43^. For the data from Camp et al. we subset the data to cells labeled “fetal” and estimated transcripts per million reads (TPM) for each gene in each cell using RSEM^63^ given the STAR^64^ mapping results. We then computed a PCA, a kNN graph, UMAP and Leiden clustering (resolution 0.2) using scanpy. We then selected the cluster with the highest STMN2 and NEUROD6 expression as the cortical neuron cluster and used only those cells. For the data from Eze et al. we subset the data sets to cells annotated as “Neuronal” in Supplementary Table 5 (“Cortex Annotations”) of their publication and computed a PCA, neighborhood graph and UMAP to visualize the data set. We found that only samples from the individuals “CS14_3”, “CS20”, “CS22” and “CS20” contained detectable expression of STMN2 and NEUROD6 so we subset the data set further to only cells from those individuals.

To compute differential expression between HNOCA cells and their primary counterparts, we first aggregated cells of the same regional neural cell type into pseudobulk samples by summing the counts for every sample (annotation columns: “batch” for HNOCA, “SampleID” for the human developing brain atlas, “sample” for Camp et al. and “individual” for Eze et al.) using the Python implementation of decoupler65 (v1.4.0) while discarding any samples with less than 10 cells or 1000 total counts. We then subsetted the feature space to the intersection of features of all data sets and removed any cells with fewer than 200 genes expressed. We further removed any genes expressed in less than 1% of neurons in HNOCA and any genes located on the X and Y chromosomes. For each regional neural cell type, we removed any sample from the pseudobulk data which was associated with an organoid differentiation assay with less than two total samples or fewer than 100 total cells. We next used edgeR^66^ to iteratively compute DE genes between each organoid differentiation protocol and primary cells of the matching regional neural cell types for every regional neural cell type while correcting for organoid age in days, number of cells per pseudobulk sample, median and standard deviation of the number of detected genes per pseudobulk sample. We used Braun et al. (the human developing brain atlas mentioned above), Eze et al. and Camp et al. data as primary data for the DE comparison in the cell type “Dorsal Telencephalic Neuron NT-VGLUT” while for all other cell types, we used the human developing brain atlas as the fetal data set. We used the edgeR genewise negative binomial generalized linear model with quasi-likelihood F-tests. We deemed a gene significantly DE if its false-discovery rate (Benjamini-Hochberg) corrected p-value was smaller than 0.05 and it had an absolute log2-fold-change above 0.5.

We used the GSEApy^67^ python package to carry out functional enrichment analysis in our DE results using the “GO_Biological_Process_2021” gene set.

### Transcriptomic similarity between HNOCA neural cell types and their primary counterparts in the human developing brain atlas

To estimate the transcriptomic similarity between neurons in HNOCA and the human developing brain atlas^12^, we first summarized the average expression of each neural cell type in the primary reference, as well as in each data set of HNOCA. For each HNOCA data set, only neural cell types with at least 20 cells were considered. Highly variable genes were identified across the neural cell types in the primary reference using a Chi-squared test-based variance ratio test on the generalized linear model with Gamma distribution (identity link), given coefficient of variance of transcript counts across neural cell types as the response, and the reciprocal of average transcript count across neural cell types as the independent variable. Genes with Benjamini-Hochberg adjusted p-value less than 0.01 were considered as highly variable genes. Similarity between one neural cell type in the primary atlas and its counterpart in each HNOCA data set was then calculated, as the Spearman correlation coefficient across the identified highly variable genes.

To estimate the similarity of the core transcriptomic identity, which is defined by the co-expression of transcription factors (TFs), the highly variable genes were subset to only TFs for calculating Spearman correlations. The list of TFs was retrieved from the AnimalTFDB 4.0 database^68^ .

To identify metabolically stressed cells in the data sets, we used the scanpy score_genes function with default parameters to score the “canonical glycolysis” gene set obtained from the enrichR “GO_Biological_Process_2021” database across all neuronal cells from HNOCA and primary references of Braun et al., Eze et al. and Camp et al.

To estimate the significance of the difference between the correlation of glycolysis scores and whole transcriptomic similarities, and the correlation of glycolysis scores and core transcriptomic identity similarities, we generated 100 subsets of highly variable genes, each with the same size as the highly variable TFs. Transcriptomic similarities were calculated based on those subsets, and then correlated with the glycolysis scores.

### Reference mapping of the neural organoid morphogen screen scRNA-seq data to the human developing brain atlas and HNOCA

We used scArches to map scRNA-seq data from the neural organoid morphogen screen to both the scANVI model of the human developing brain atlas^12^ and the scPoli model of the HNOCA. In both cases, the “dataset” field of the screen data was used as the batch covariate, which indicates belonging to either of the three categories: “organoid screen”, “secondary organoid screen”, or “fetal striatum 21pcw”. For mapping to the primary reference, we used the scvi-tools implementation of scArches without the use of cell type annotations and trained the model for 500 epochs with weight_decay = 0 and otherwise default parameters. For mapping to HNOCA we used scArches through scPoli and trained the model for 500 epochs without unlabeled prototype training.

## Data and code availability

All curated individual HNOCA datasets are available for easy access via the sfaira python tool48. The integrated HNOCA data will be uploaded to Zenodo and cellxgene. A Shinyapp is in preparation for online exploration of the HNOCA data. The snapseed package is available at https://github.com/devsystemslab/snapseed. Jupyter notebooks to reproduce the analysis are available at https://github.com/theislab/neural_organoid_atlas.

## ACKNOWLEDGEMENTS

We thank C. DeDonno for his support in improving our data integration efforts using scPoli. We thank D. Klein, P. Weiler and M. Lange for insightful discussions on the moscot framework, (neural) optimal transport and real time informed pseudo-time analyses. We thank F. Sanchis-Calleja, S. Jansen and F. Zenk for insightful comments on summarizing neural organoid protocols. We thank P. Lönnerberg and S. Linnarsson for insightful discussions on the application of the human developing brain atlas in this study. We thank A. Fiorenzano, A.R. Kriegstein, A.R. Muotri, B.G. Novitch, G. Ming, I. Park, J.A. Knoblich, M.A. Lancaster, P. Arlotta, S. Temple, S.P. Paşca, T. Kato for their support on data and metadata retrieval. This work was supported by Chan Zuckerberg Initiative DAF, an advised fund of the Silicon Valley Community Foundation (CZF2019-002440 and CZF2021-237566, to J.G.C. and B.T.). This work was co-funded by the Swiss National Science Foundation (project grant 310030_192604, to B.T.). This work was co-funded by the European Union (ERC, DeepCell -101054957, to A.S. and F.J.T.; ERC, Organomics -758877, to B.T.; H2020, Braintime -874606, to B.T.; ERC, Anthropoid -803441, to J.G.C.). Views and opinions expressed are however those of the authors only and do not necessarily reflect those of the European Union or the European Research Council. Neither the European Union nor the granting authority can be held responsible for them. This work was supported by the BMBF-funded de.NBI Cloud within the German Network for Bioinformatics Infrastructure (de.NBI) (031A532B, 031A533A, 031A533B, 031A534A, 031A535A, 031A537A, 031A537B, 031A537C, 031A537D, 031A538A) (to L.D., A.S., K.X.L. and I.S.). This work was supported through a Fulbright grant of the German-American Fulbright Commission (to K.X.L.). L.D. acknowledges support by the Joachim Herz Foundation. This publication is part of the Human Cell Atlas (www.humancellatlas.org/publications/).

## AUTHOR CONTRIBUTIONS

Z.H., L.D., J.S. collected and retrieved the scRNA-seq data involved in HNOCA, with suggestions from S.P.P., J.G.C. and B.T.. H.L. and M.S. generated the unpublished midbrain organoid data. A.A and G.Q. generated and shared the unpublished cerebellar organoid data. J.S.F. developed snapseed. Z.H. curated cell type hierarchy with the support from J.S.F. and L.D.. L.D., K.X.L., I.S. and A.S. performed HNOCA data curation and metadata harmonization. L.D., with the support from K.X.L. and I.S., performed HNOCA data preprocessing and integration using the pipeline developed by Z.H., L.D., J.S.F. and A.S.. L.D. and K.X.L. performed the benchmark of integration methods. Z.H. did HNOCA cell type annotation. K.X.L. and J.S.F. performed the real-time-informed pseudotime analysis. J.S.F. performed reference mapping of HNOCA to the human developing brain atlas with support from A.S.. Z.H. developed and performed label transfer and presence score estimation. I.S. and J.S.F. performed morphogen analysis, with the organoid protocols summarized by Z.H.. L.D. and Z.H. with support from I.S. and K.X.L. performed DE and transcriptomic comparison analysis. J.S.F. and A.S. performed reference mapping of organoid morphogen screen data set to HNOCA and the human developing brain atlas and the followup analysis. Z.H., J.G.C., F.J.T. and B.T. designed the project. Z.H., L.D., J.S.F., A.S., I.S., S.P.P., J.G.C., F.J.T. and B.T. wrote the manuscript with input from all the coauthors. All authors read and approved the final manuscript.

## Extended Data Figures

**Extended Data Fig. 1.**
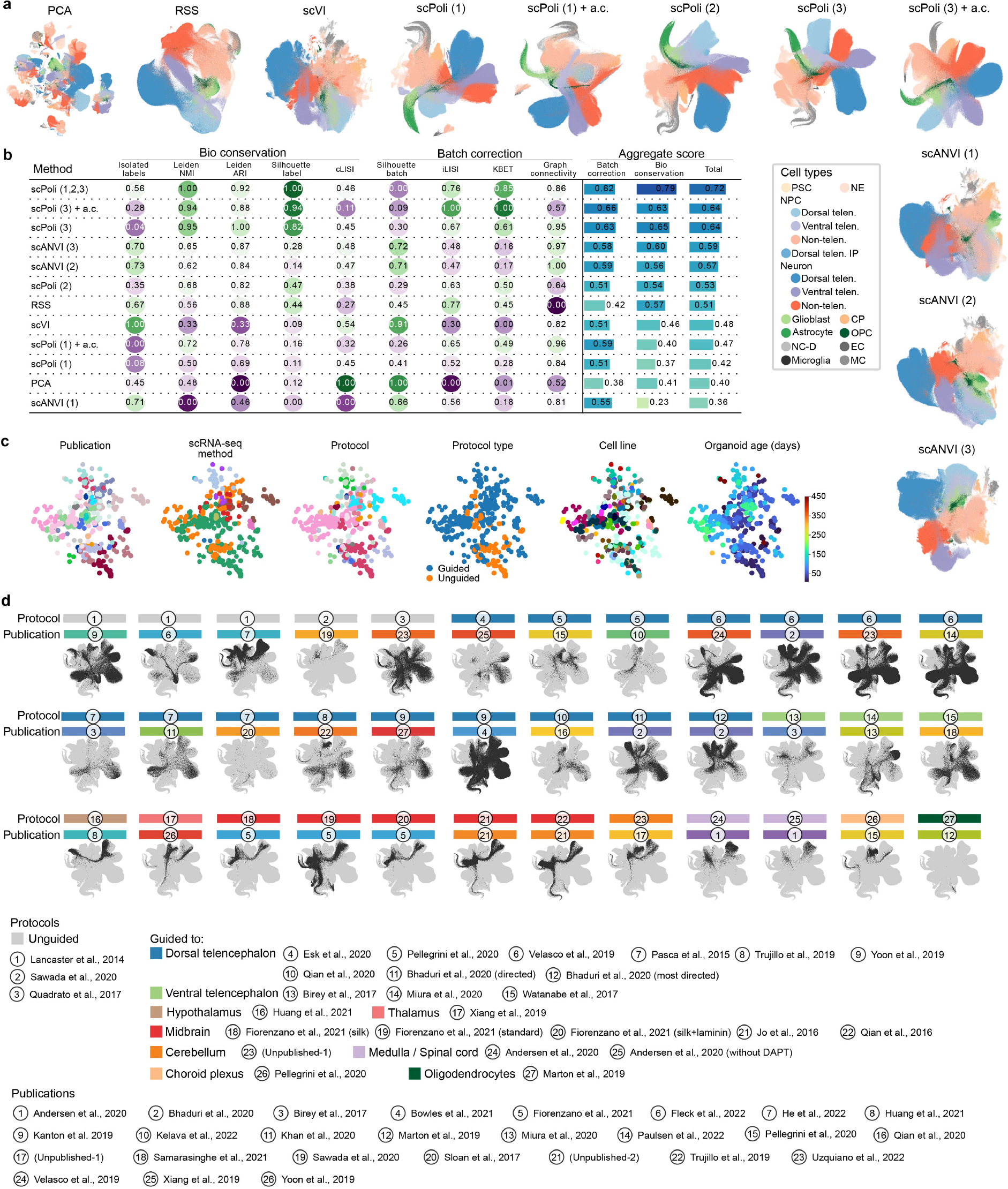
Benchmark of data integration. (a) UMAPs of HNOCA, either without any data integration (PCA) or with different data integration methods applied. Number in parenthesis indicates which level of RSS-based snapseed annotation labels were provided as input to the model for methods which support semi-supervised data integration. Dots in all UMAP embeddings, each of which represents a cell, are colored by the cell type annotation introduced in Figure 1. a.c. = aggrecell algorithm (b) scIB benchmarking metrics on all tested integration methods. (c) PCA of the scPoli sample embeddings from the final scPoli integration of HNOCA presented throughout the manuscript, colored by publications, scRNA-seq methods, organoid protocols, protocol types, cell lines, and sample ages. (d) UMAPs of HNOCA based on the final scPoli integration, each with one data set highlighted. Here, one data set is defined as data representing one protocol in one publication. The protocol and publication of each data set are shown by the color bar and indices on top of the UMAP.

**Extended Data Fig. 2.**
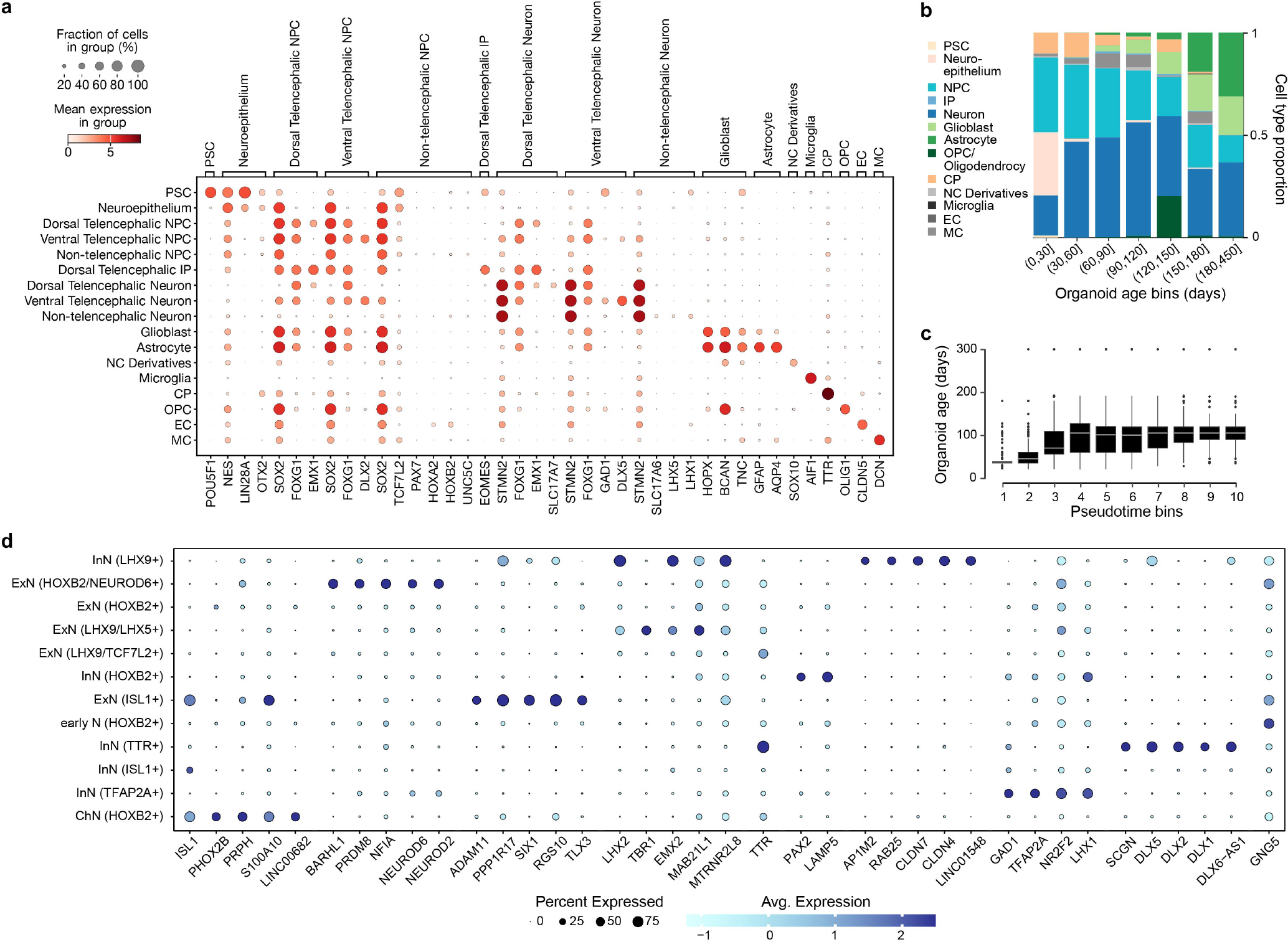
Characterization of HNOCA. (a) Expression of selected marker genes used in the semi-automatic annotation of cell types for Figure 1. (b) Mean cell type proportion over all data sets per organoid age bin. (c) Distribution of sample real-time age in days over deciles of computed pseudotime. (d) Expression of top markers in different non-telencephalic neural cell types. Markers are defined as genes with AUC>0.7, in-out detection rate difference>20%, in-out detection rate ratio>2 and fold change>1.2. When more than 5 markers are found, only the top-5 (with the highest in-out detection rate ratio) are shown.

**Extended Data Fig. 3.**
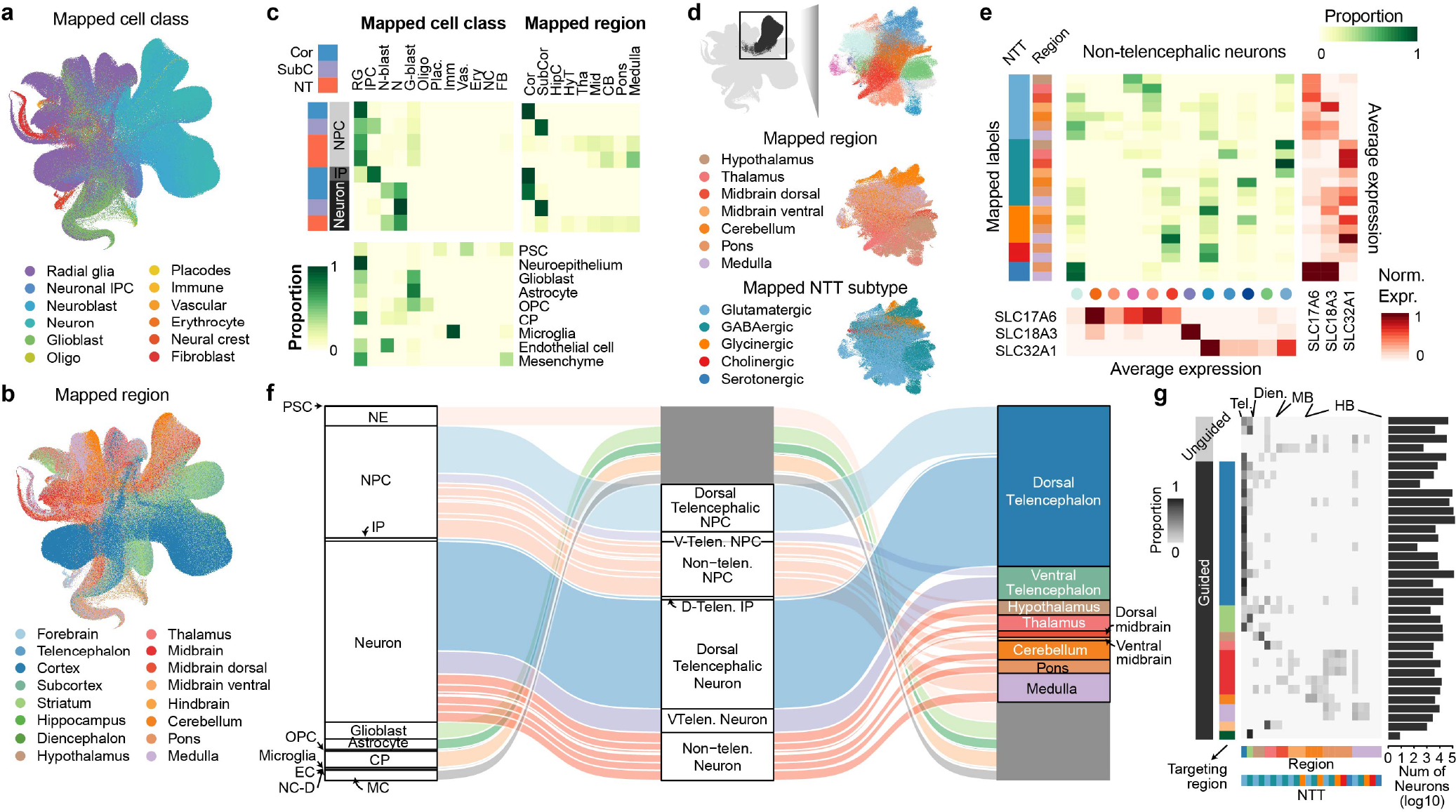
Mapping-assisted annotation refinement of HNOCA. (a-b) UMAP of HNOCA colored by the mapped (a) cell classes and (b) brain regions, both from the human developing brain cell atlas^12^ as the primary reference. (c) Comparison of the HNOCA cell type annotation with the primary reference mapping-based transferred cell class and brain region labels. Darkness of cells indicates proportions of each HNOCA cell type being assigned to different cell class and brain region categories. Brain region labels are only shown for the HNOCA neural cell types. (d) UMAP of non-telencephalic neurons, colored by clusters (upper), mapped brain regions (middle) and mapped neurotransmitter transporter (NTT) subtypes (bottom). (e) Comparison of non-telencephalic neural cell types, defined as the concatenation of the mapped brain region and NTT subtype, with the clusters. The middle heatmap shows contributions of different clusters to different neural cell types. The sidebar on the left shows the neural cell types; dots under the heatmap show clusters. The heatmaps on the bottom and on the right show the average expression of three neurotransmitter transporters SLC17A6, SLC18A3 and SLC32A1 in clusters (bottom) and neural cell types (right). (f) Overview of the HNOCA cell type composition for the first two levels of the cell annotation (left -level-1, middle -level-2), and the refined regional annotation assisted by mapping of non-telencephalic NPC and neurons to the primary reference (right). (g) neural cell type compositions of different data sets (rows). Darkness of the heatmap shows the proportions of different neural cell types per HNOCA data set. Sidebars on the left show organoid protocol types of different data sets. Sidebars on the bottom show neural cell types. Bars on the right show total neuron numbers across data sets.

**Extended Data Fig. 4.**
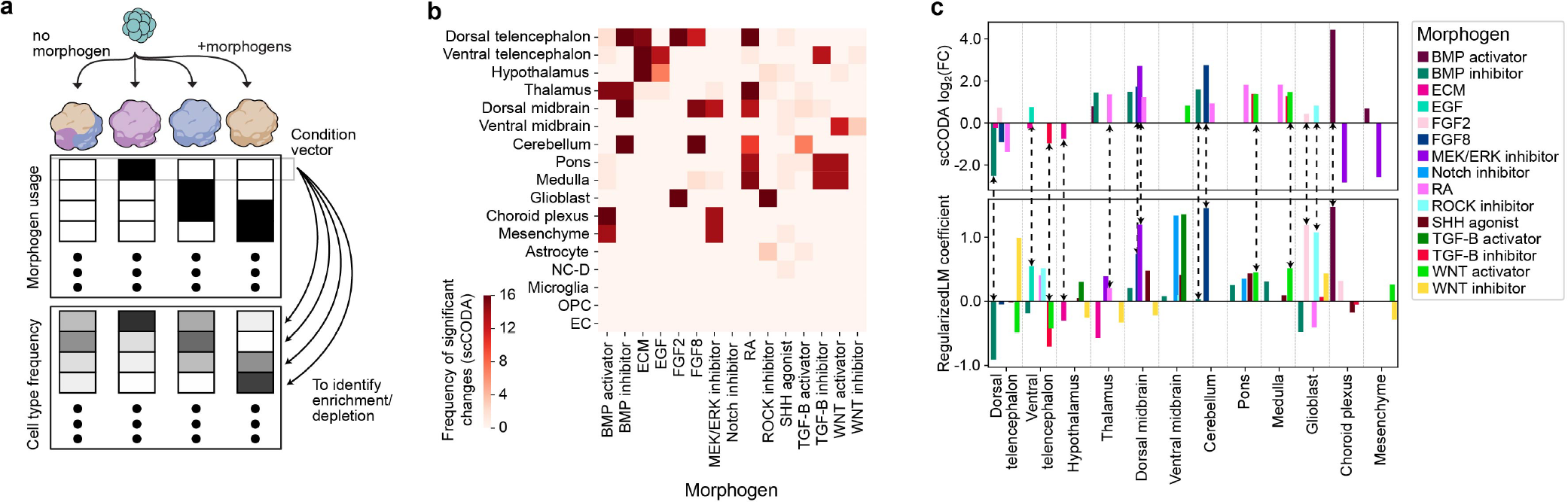
Relationship between morphogen usage and cell type as well as regional composition. (a) Schematic of estimating cell type enrichment with different morphogen usages. (b) This heat map indicates in how many of the 17 iterations sc-CODA was executed (using each of the 17 regional cell identity as a reference once) the respective morphogen was found to lead to compositional changes with respect to the reference regional cell identity. A morphogen effect was called significant in this consensus approach if it had a significant effect on cell type composition with respect to more than half of the reference cell types. (c) Effect of different morphogens on regional organoid composition in HNOCA. Positive values correspond to a higher abundance of cells from the indicated regional cell identity in cases where the respective morphogen was used in the differentiation protocol. Top: log2-fold-effect sizes of morphogens per regional cell identity as computed by the scCODA model. Bottom: L1-regularized linear model coefficients. The dashed arrows show consistent enrichment/depletion identified by the two methods.

**Extended Data Fig. 5.**
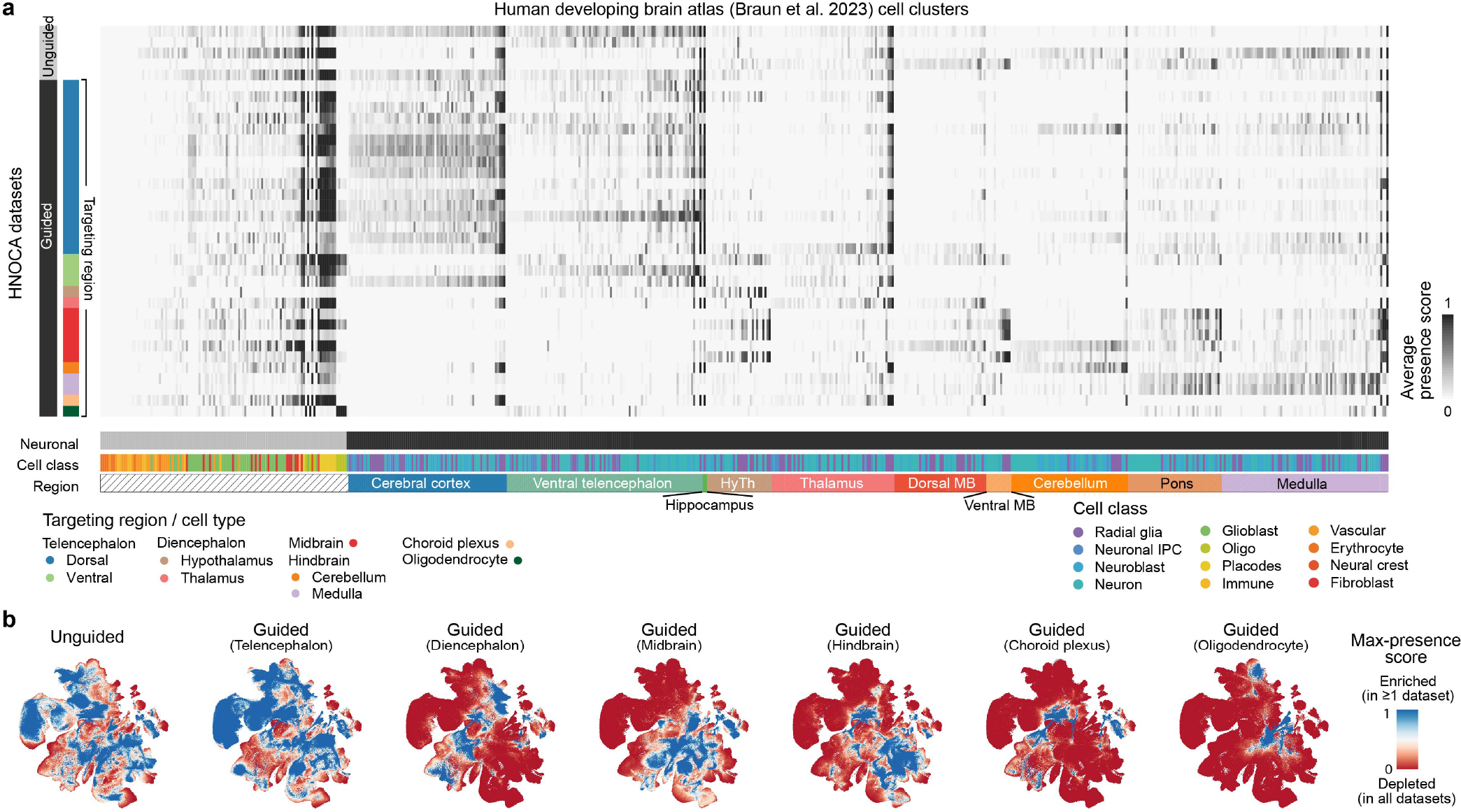
Presence scores per HNOCA data set. (a) Average normalized presence scores of different HNOCA data sets (rows) in different cell clusters in the primary reference of the human developing brain atlas12 (columns). Sidebars on the left show organoid differentiation protocol types of HNOCA data sets. Sidebars underneath show cell class and the commonest region information of the cell clusters in the primary reference (HyTh -hypothalamus, MB -midbrain). (b) UMAP of the primary reference, colored by the max presence scores across different HNOCA data subsets, split by organoid protocol types. A high max presence score suggests enrichment of the corresponding primary cell state in at least one HNOCA data set among the data sets based on the specific type of organoid protocols, with a low score meaning under-representation of the cell state in all data sets in the subset.

**Extended Data Fig. 6.**
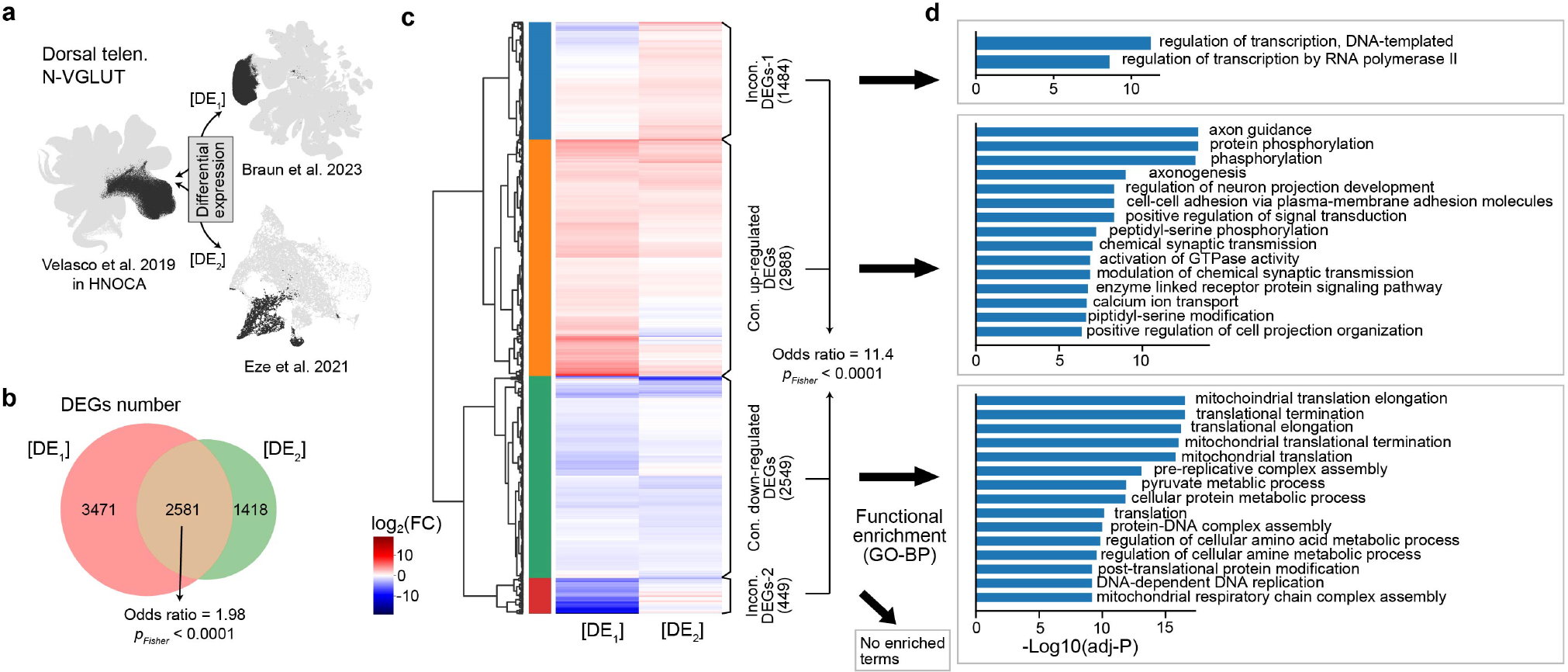
Robustness of organoid-primary DEGs against primary reference, and diseases-associated organoid-primary DEGs. (a) Schematic of DE analysis between a subset of dorsal telencephalic neurons in HNOCA (Velasco et al. 2019^19^) with counterparts in two developing human brain atlases^12,43^. (b) Number of DEGs between cortical neurons from Velasco et al. organoid data and primary fetal cortical neurons from Braun et al. 2023 (10x 3’ v2 chemistry only, [DE1]) or Eze et al. 2021 ([DE2]) respectively. (c) Log2-transformed fold changes (log2FC) across all 7470 DEGs between dorsal telencephalic neurons from Velasco et al. and either primary fetal cortical neurons from Braun et al. (10x 3’ v2 chemistry only) or Eze et al.. The dendrogram shows the hierarchical clustering of DEGs based on their log2FC against the two primary data. (d) Functional enrichment analysis of the four clusters of DEGs. Widths of bars show -log10-transformed adjusted P-value of Fisher’s exact test. Only the first 15 terms with adjusted P<0.05 are shown.

**Extended Data Fig. 7.**
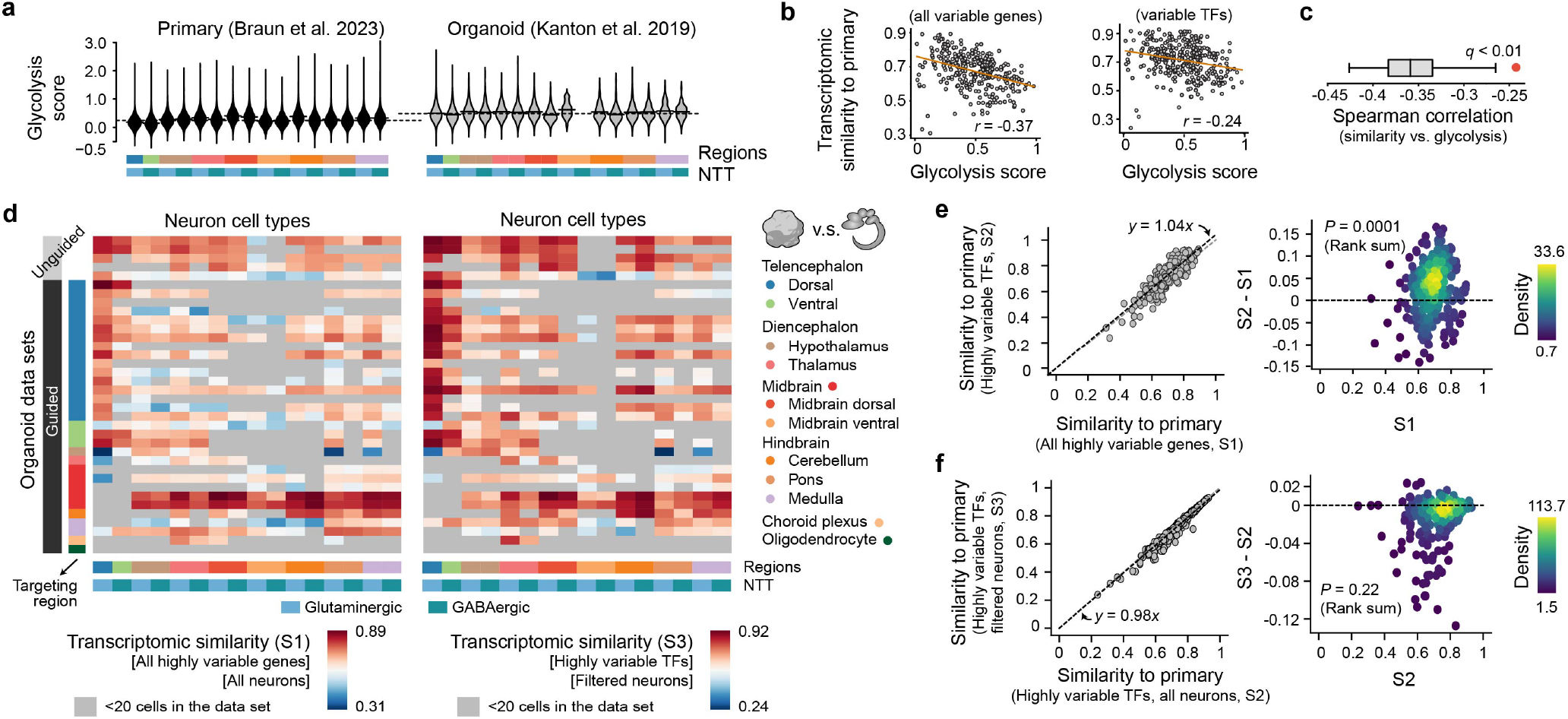
Transcriptomic fidelity of neurons and cell stress. (a) Glycolysis scores of different neural cell types in primary (left, Braun et al.^12^) and a selected organoid data set (right, Kanton et al.^16^). (b) Correlation between average glycolysis scores and transcriptomic similarities (Spearman correlation) to primary counterparts. Each dot represents one neural cell type generated by one protocol. The correlation is calculated based on either all variable genes (left) or variable transcription factors (TFs, right). (c) The correlation between glycolysis scores and transcriptomic similarities to primary is significantly weaker when only TFs are taken into consideration. The box shows the distribution of correlation when a random subset of variable genes, with the same number as the variable TFs, are used. The red dot shows the correlation using variable TFs. (d) Spearman correlation between average gene expression profiles of neural cell types in HNOCA and those in the primary reference of human developing brain atlas^12^, across either all the variable genes (left, *S*_1_) or variable transcriptional factors (right, *S*_3_). The average gene expression profile per neural cell type was calculated with all cells (*S*_1_) or cells with low glycolysis scores (glycolysis score < 0.6, *S*_3_). (e) Core transcriptomic fidelity of organoid neurons (*S*_2_, shown in Fig.3) which only considers TFs, is higher than the global transcriptomic fidelity (*S*_1_) which considers all the highly variable genes. Core transcriptomic fidelity and global transcriptomic fidelity are highly correlated (left, x-axis -*S*_1_, y-axis -*S*_2_, each dot represents one neural cell type in one HNOCA data set), while core transcriptomic fidelity is significantly higher (right, x-axis: *S*_1_, y-axis: *S*_2_ *−S*_1_, dots are colored by density estimated with Gaussian kernel). (f) Core transcriptomic fidelity of organoid neurons is not improved by filtering cells based on glycolysis scores. The left panel shows core transcriptomic fidelities without (x-axis, *S*_2_) and with glycolysis score filtering (glycolysis score<0.6, y-axis, *S*_3_). The Wilcoxon test suggests *S*_2_ and *S*_3_ are not significantly different (right, x-axis: *S*_2_, y-axis: *S*_3_ *−S*_2_, dots are colored by density estimated with Gaussian kernel).

**Extended Data Fig. 8.**
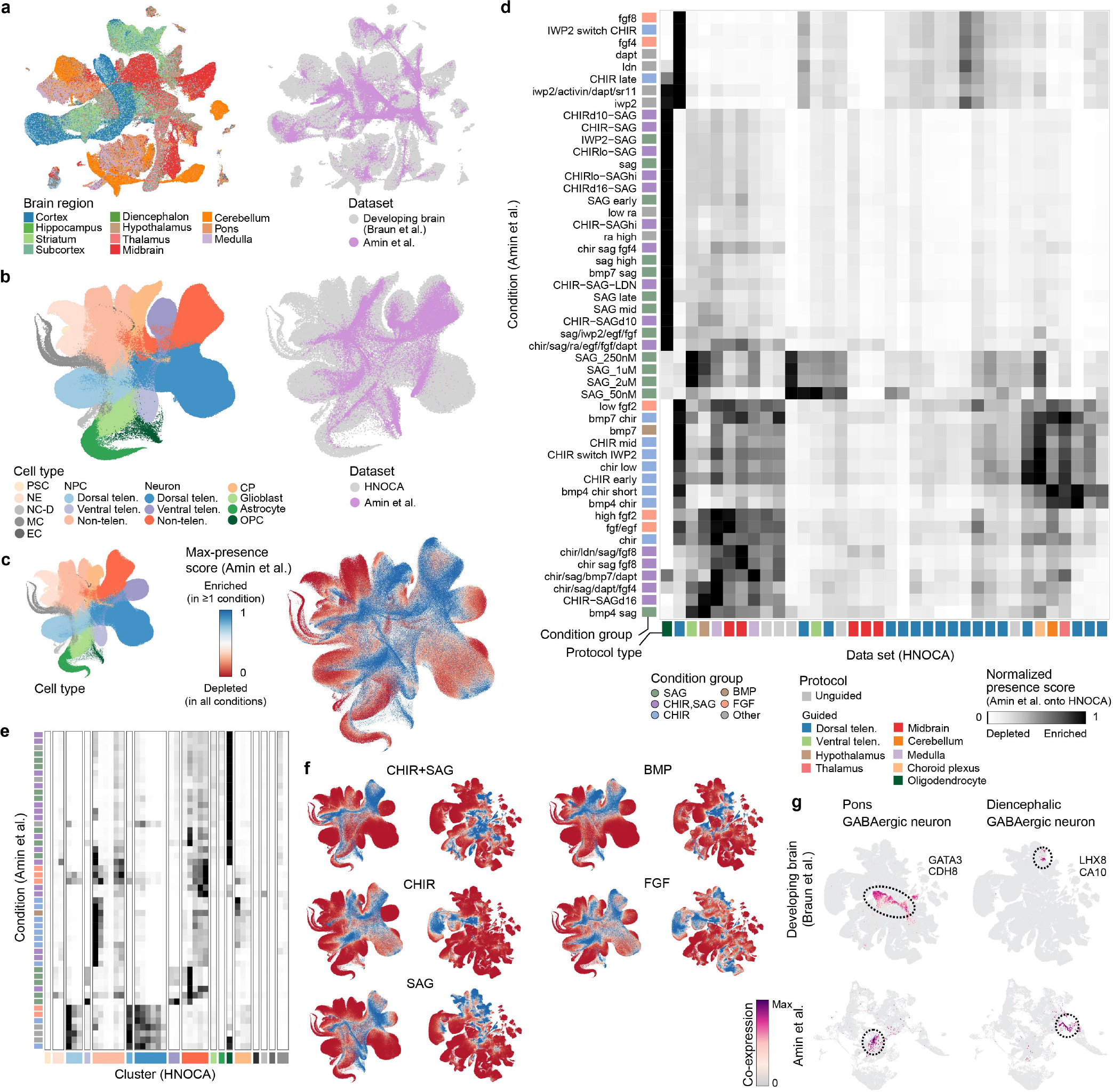
Reference mapping of the neural organoid morphogen screen data to HNOCA and the human developing brain atlas. (a) UMAP embedding of the human developing brain atlas and neural organoid morphogen screen^13^ data sets based on the joint scANVI latent space colored by brain region (left) and data set (right). (b) UMAP embedding of HNOCA and the screen data sets based on the joint scPoli latent space colored by annotated cell type (left) and data set (right). (c) scPoli UMAP embedding of the HNOCA colored by cell type (left) and max presence score across all data sets (right). (d) Heatmap showing min-max scaled average presence scores of each condition in the screen data set in HNOCA data sets. (e) Heatmap showing min-max scaled average presence scores of each condition in the screen data set in each leiden cluster in HNOCA, ordered by annotated cell type. (f) UMAP embeddings of HNOCA (left) and the human developing brain atlas (right) colored by presence scores for each condition group in the screen data set. (g) UMAP embeddings of the human developing brain atlas (upper) and screen data set (lower) colored by coexpression scores of clusters with gained coverage in the screen data set.

## REFERENCES

1 Velasco, S., Paulsen, B. Arlotta, P. 3D Brain Organoids: Studying Brain Development and Disease Outside the Embryo. Annu. Rev. Neurosci. 43, 375–389 (2020).

2 Sidhaye, J. Knoblich, J. A. Brain organoids: an ensemble of bioassays to investigate human neurodevelopment and disease. Cell Death Differ. 28, 52–67 (2020).

3 Paşca, S. P. et al. A nomenclature consensus for nervous system organoids and assembloids. Nature 609, 907–910 (2022).

4 Paşca, A. M. et al. Functional cortical neurons and astrocytes from human pluripotent stem cells in 3D culture. Nat. Methods 12, 671–678 (2015).

5 Xiang, Y. et al. hESC-Derived Thalamic Organoids Form Reciprocal Projections When Fused with Cortical Organoids. Cell Stem Cell 24, 487–497.e7 (2019).

6 Miura, Y. et al. Generation of human striatal organoids and cortico-striatal assembloids from human pluripotent stem cells. Nat. Biotechnol. 38, 1421–1430 (2020).

7 Huang, W.-K. et al. Generation of hypothalamic arcuate organoids from human induced pluripotent stem cells. Cell Stem Cell 28, 1657–1670.e10 (2021).

8 Jo, J. et al. Midbrain-like Organoids from Human Pluripotent Stem Cells Contain Functional Dopaminergic and Neuromelanin-Producing Neurons. Cell Stem Cell 19, 248–257 (2016).

9 Lancaster, M. A. et al. Cerebral organoids model human brain development and microcephaly. Nature 501, 373–379 (2013).

10 Quadrato, G. et al. Cell diversity and network dynamics in photosensitive human brain organoids. Nature 545, 48–53 (2017).

11 Camp, J. G. et al. Human cerebral organoids recapitulate gene expression programs of fetal neocortex development. Proc. Natl. Acad. Sci. U. S. A. 112, 15672–15677 (2015).

12 Braun, E. et al. Comprehensive cell atlas of the first-trimester developing human brain. bioRxiv 2022.10.24.513487 (2022) doi:10.1101/2022.10.24.513487.

13 Amin, N. D. et al. Generating human neural diversity with a multi plexed morphogen screen in organoids. bioRxiv 2023.05.31.541819 (2023) doi:10.1101/2023.05.31.541819.

14 Birey, F. et al. Assembly of functionally integrated human forebrain spheroids. Nature 545, 54–59 (2017).

15 Sloan, S. A. et al. Human Astrocyte Maturation Captured in 3D Cerebral Cortical Spheroids Derived from Pluripotent Stem Cells. Neuron 95, 779–790.e6 (2017).

16 Kanton, S. et al. Organoid single-cell genomic atlas uncovers human-specific features of brain development. Nature 574, 418–422 (2019).

17 Marton, R. M. et al. Differentiation and maturation of oligodendrocytes in human three-dimensional neural cultures. Nat. Neurosci. 22, 484–491 (2019).

18 Trujillo, C. A. et al. Complex Oscillatory Waves Emerging from Cortical Organoids Model Early Human Brain Network Development. Cell Stem Cell 25, 558–569.e7 (2019).

19 Velasco, S. et al. Individual brain organoids reproducibly form cell diversity of the human cerebral cortex. Nature 570, 523–527 (2019).

20 Yoon, S.-J. et al. Reliability of human cortical organoid generation. Nat. Methods 16, 75–78 (2019).

21 Andersen, J. et al. Generation of Functional Human 3D Cortico-Motor Assembloids. Cell 183, 1913–1929.e26 (2020).

22 Bhaduri, A. et al. Cell stress in cortical organoids impairs molecular subtype specification. Nature 578, 142–148 (2020).

23 Pellegrini, L. et al. Human CNS barrier-forming organoids with cerebrospinal fluid production. Science 369, (2020).

24 Qian, X. et al. Sliced Human Cortical Organoids for Modeling Distinct Cortical Layer Formation. Cell Stem Cell 26, 766–781.e9 (2020).

25 Sawada, T. et al. Developmental excitation-inhibition imbalance underlying psychoses revealed by single-cell analyses of discordant twins-derived cerebral organoids. Mol. Psychiatry 25, 2695–2711 (2020).

26 Khan, T. A. et al. Neuronal defects in a human cellular model of 22q11.2 deletion syndrome. Nat. Med. 26, 1888–1898 (2020).

27 Bowles, K. R. et al. ELAVL4, splicing, and glutamatergic dysfunction precede neuron loss in MAPT mutation cerebral organoids. Cell 184, 4547–4563.e17 (2021).

28 Fiorenzano, A. et al. Single-cell transcriptomics captures features of human midbrain development and dopamine neuron diversity in brain organoids. Nat. Commun. 12, 7302 (2021).

29 Samarasinghe, R. A. et al. Identification of neural oscillations and epileptiform changes in human brain organoids. Nat. Neurosci. 24, 1488–1500 (2021).

30 Fleck, J. S. et al. Inferring and perturbing cell fate regulomes in human brain organoids. Nature (2022) doi:10.1038/s41586-022-05279-8.

31 He, Z. et al. Lineage recording in human cerebral organoids. Nat. Methods 19, 90–99 (2022).

32 Kelava, I., Chiaradia, I., Pellegrini, L., Kalinka, A. T. Lancaster, M. A. Androgens increase excitatory neurogenic potential in human brain organoids. Nature 602, 112–116 (2022).

33 Uzquiano, A. et al. Proper acquisition of cell class identity in organoids allows definition of fate specification programs of the human cerebral cortex. Cell 185, 3770–3788.e27 (2022).

34 Vértesy, Á. et al. Gruffi: an algorithm for computational removal of stressed cells from brain organoid transcriptomic datasets. EMBO J. 41, e111118 (2022).

35 Paulsen, B. et al. Autism genes converge on asynchronous development of shared neuron classes. Nature 602, 268–273 (2022).

36 De Donno, C. et al. Population-level integration of single-cell datasets enables multi-scale analysis across samples. bioRxiv 2022.11.28.517803 (2022) doi:10.1101/2022.11.28.517803.

37 Luecken, M. D. et al. Benchmarking atlas-level data integration in single-cell genomics. Nat. Methods 19, 41–50 (2022).

38 Traag, V. A., Waltman, L. van Eck, N. J. From Louvain to Leiden: guaranteeing well-connected communities. Sci. Rep. 9, 5233 (2019).

39 Klein, D. et al. Mapping cells through time and space with moscot. bioRxiv 2023.05.11.540374 (2023) doi:10.1101/2023.05.11.540374.

40 Lopez, R., Regier, J., Cole, M. B., Jordan, M. I. Yosef, N. Deep generative modeling for single-cell transcriptomics. Nat. Methods 15, 1053–1058 (2018).

41 Xu, C. et al. Probabilistic harmonization and annotation of single-cell transcriptomics data with deep generative models. Mol. Syst. Biol. 17, e9620 (2021).

42 Lotfollahi, M. et al. Query to reference single-cell integration with transfer learning. bioRxiv 2020.07.16.205997 (2020) doi:10.1101/2020.07.16.205997.

43 Eze, U. C., Bhaduri, A., Haeussler, M., Nowakowski, T. J. Kriegstein, A. R. Single-cell atlas of early human brain development highlights heterogeneity of human neuroepithelial cells and early radial glia. Nat. Neurosci. 24, 584–594 (2021).

44 Ashburner, M. et al. Gene Ontology: tool for the unification of biology. Nat. Genet. 25, 25–29 (2000).

45 Aleksander, S. A. et al. The Gene Ontology knowledgebase in 2023. Genetics 224, iyad031 (2023).

46 Qian, X., Song, H. Ming, G.-L. Brain organoids: advances, applications and challenges. Development 146, (2019).

47 Stern, C. D. et al. Head-tail patterning of the vertebrate embryo: one, two or many unresolved problems? Int. J. Dev. Biol. 50, 3–15 (2006).

48 Fischer, D. S. et al. Sfaira accelerates data and model reuse in single cell genomics. Genome Biol. 22, 248 (2021).

49 Cunningham, F. et al. Ensembl 2022. Nucleic Acids Res. 50, D988–D995 (2022).

50 Virshup, I., Rybakov, S., Theis, F. J., Angerer, P. Wolf, F. A. anndata: Annotated data. bioRxiv (2021) doi:10.1101/2021.12.16.473007.

51 Wolf, F. A., Angerer, P. Theis, F. J. SCANPY: large-scale single-cell gene expression data analysis. Genome Biol. 19, 15 (2018).

52 McInnes, L., Healy, J., Saul, N. Großberger, L. UMAP: Uniform Manifold Approximation and Projection. J. Open Source Softw. 3, 861 (2018).

53 Nolet, C. et al. Accelerating single-cell genomic analysis with GPUs. bioRxiv 2022.05.26.493607 (2022) doi:10.1101/2022.05.26.493607.

54 Lotfollahi, M. et al. Mapping single-cell data to reference atlases by transfer learning. Nat. Biotechnol. 40, 121–130 (2022).

55 GitHub - YosefLab/scib-metrics: Accelerated, Python-only, single-cell integration benchmarking metrics. GitHub https://github.com/YosefLab/scib-metrics.

56 Lange, M. et al. CellRank for directed single-cell fate mapping. Nat. Methods 19, 159–170 (2022).

57 Haghverdi, L., Buettner, F. Theis, F. J. Diffusion maps for high-dimensional single-cell analysis of differentiation data. Bioinformatics 31, 2989–2998 (2015).

58 Gayoso, A. et al. A Python library for probabilistic analysis of single-cell omics data. Nat. Biotechnol. 40, 163–166 (2022).

59 Virshup, I. et al. The scverse project provides a computational ecosystem for single-cell omics data analysis. Nat. Biotechnol. 41, 604–606 (2023).

60 GitHub - theislab/pertpy: Perturbation Analysis in the scverse ecosystem. GitHub https://github.com/theislab/pertpy.

61 Büttner, M., Ostner, J., Müller, C. L., Theis, F. J. Schubert, B. scCODA is a Bayesian model for compositional single-cell data analysis. Nat. Commun. 12, 1–10 (2021).

62 Tay, J. K., Narasimhan, B. Hastie, T. Elastic Net Regularization Paths for All Generalized Linear Models. J. Stat. Softw. 106, (2023).

63 Li, B. Dewey, C. N. RSEM: accurate transcript quantification from RNA-Seq data with or without a reference genome. BMC Bioinformatics 12, 323 (2011).

64 Dobin, A. et al. STAR: ultrafast universal RNA-seq aligner. Bioinformatics 29, 15–21 (2013).

65 Badia-I-Mompel, P. et al. decoupleR: ensemble of computational methods to infer biological activities from omics data. Bioinform Adv 2, vbac016 (2022).

66 Robinson, M. D., McCarthy, D. J. Smyth, G. K. edgeR: a Bioconductor package for differential expression analysis of digital gene expression data. Bioinformatics 26, 139–140 (2010).

67 Fang, Z., Liu, X. Peltz, G. GSEApy: a comprehensive package for performing gene set enrichment analysis in Python. Bioinformatics 39, (2023).

68 Shen, W.-K. et al. AnimalTFDB 4.0: a comprehensive animal transcription factor database updated with variation and expression annotations. Nucleic Acids Res. 51, D39–D45 (2023).

