## Supplementary Data 1 for "An integrated transcriptomic cell atlas of human neural organoids": Data_S1_snapseed_markers.yaml

```
# Brain cell types and markers
neural_progenitor_cell:
  marker_genes: [SOX2, VIM, NES]
  subtypes: 
    glioblast:
      marker_genes: [HOPX, BCAN, TNC] 
    telencephalic_npc:
      marker_genes: [FOXG1]
      subtypes: 
        dorsal_npc:
          marker_genes: [EMX1]
        hippocampal_npc:
          marker_genes: [GABRA2, GABRB1] 
        ventral_npc:
          marker_genes: [DLX2]
    diencephalic_npc:
      marker_genes: [TCF7L2, SIX3]
      subtypes: 
        hypothalamic_npc:
          marker_genes: [SIX3]
        thalamic_npc:
          marker_genes: [TCF7L2]
    mesencephalic_npc:
      marker_genes: [OTX2, PAX7]
    retinal_npc:
      marker_genes: [VSX2]
    rhombencephalic_npc:
      marker_genes: [CYP26A1, HOXA2, HOXB2, UNC5C, BCAN] 
      subtypes: 
        cerebellar_npc:
          marker_genes: [UNC5C, CYP26A1]
        pons_npc:
          marker_genes: [GPC6, BCAN]
        medullary_npc:
          marker_genes: [HOXB2, HOXD3, HOXA3]
neuron:
  marker_genes: [STMN2, DCX]
  subtypes:
    excitatory_neuron:
      marker_genes: [SLC17A7, SLC17A6]
      subtypes:
        telencephalic_excitatory_neuron:
          marker_genes: [FOXG1, EMX1, SLC17A7]
          subtypes:
            deeper_layer_cortical_excitatory_neuron:
              marker_genes: [BCL11B]
            upper_layer_cortical_excitatory_neuron:
              marker_genes: [SATB2]
            hippocampal_excitatory_neuron:
              marker_genes: [GABRA2, GABRB1]
        non_telencephalic_excitatory_neuron:
          marker_genes: [SLC17A6]
          subtypes:
            diencephalic_excitatory_neuron:
              marker_genes: [TCF7L2, PITX2]
              subtypes: 
                hypothalamic_excitatory_neuron:
                  marker_genes: [PITX2]
                thalamic_excitatory_neuron:
                  marker_genes: [TCF7L2]
            mesencephalic_excitatory_neuron:
              marker_genes: [BARHL2, TFAP2D]
            rhombencephalic_excitatory_neuron:
              marker_genes: [TFAP2A, HOXA2, HOXB2, HOXB2, HOXD3, HOXA3]
              subtypes:
                cerebellar_excitatory_neuron:
                  marker_genes: [UNCX, INSM1, UNC5C]
                pons_excitatory_neuron:
                  marker_genes: [ROBO1, NEGR1]
                medullary_excitatory_neuron:
                  marker_genes: [HOXB2, HOXD3, HOXA3]
    inhibitory_neuron:
      marker_genes: [GAD1, GAD2, SLC32A1]
      subtypes:
        telencephalic_inhibitory_neuron:
          marker_genes: [FOXG1]
          subtypes:
            ventral_inhibitory_neuron:
              marker_genes: [DLX2, DLX5]
              subtypes:
                mge_inhibitory_neuron:
                  marker_genes: [NKX2-1]
                lge_inhibitory_neuron:
                  marker_genes: [ISL1]
                cge_inhibitory_neuron:
                  marker_genes: [NR2F1]
        non_telencephalic_inhibitory_neuron:
          marker_genes: [LHX5, LHX1] 
          subtypes:
            diencephalic_inhibitory_neuron:
              marker_genes: [TCF7L2, ISL1]
              subtypes:
                hypothalamic_inhibitory_neuron:
                  marker_genes: [ISL1]
                thalamic_inhibitory_neuron:
                  marker_genes: [DLX1, DLX5]
            mesencephalic_inhibitory_neuron:
              marker_genes: [OTX2]
            rhombencephalic_inhibitory_neuron:
              marker_genes: [SKOR2, HOXA2, HOXB2, CA8, LAMP5]
              subtypes:
                cerebellar_inhibitory_neuron:
                  marker_genes: [CA8, TFAP2A, SKOR2, UNC5C]
                medullary_inhibitory_neuron:
                  marker_genes: [HOXB2, HOXD3, HOXA3, LAMP5]
                pons_inhibitory_neuron:
                  marker_genes: [ROBO1, NEGR1]
choroid_plexus_epithelium:
  marker_genes: [TTR]
astrocyte:
  marker_genes: [GFAP, AQP4]
oligodendrocyte_lineage:
  marker_genes: [OLIG1, MBP]
  subtypes:
    oligodendrocyte_precursor_cell:
      marker_genes: [OLIG1]
    mature_oligodendrocyte:
      marker_genes: [MBP]
microglia:
  marker_genes: [AIF1]
vascular_endothelial_cell:
  marker_genes: [CLDN5]
mesenchymal_cell:
  marker_genes: [DCN]
neural_crest:
  marker_genes: [SOX10]
pns_neurons:
  marker_genes: [PRPH]


# All markers in one unqiue list 
# all_markers: ['SOX2', 'VIM', 'NES', 'FOXG1', 'EMX1', 'HOPX', 'GABRA2', 'GABRB1', 'DLX2', 'TCF7L2', 'SIX3', 'OTX2', 'PAX7', 'VSX2', 'CYP26A1', 'HOXA2', 'HOXB2', 'UNC5C', 'BCAN', 'GPC6', 'HOXD3', 'HOXA3', 'STMN2', 'DCX', 'SLC17A7', 'SLC17A6', 'BCL11B', 'SATB2', 'PITX2', 'BARHL2', 'TFAP2D', 'TFAP2A', 'UNCX', 'INSM1', 'ROBO1', 'NEGR1', 'GAD1', 'GAD2', 'SLC32A1', 'DLX5', 'NKX2-1', 'ISL1', 'NR2F1', 'LHX5', 'LHX1', 'CA8', 'LAMP5', 'SKOR2', 'AQP4', 'GFAP', 'TTR', 'OLIG1', 'MBP', 'AIF1', 'CLDN5', 'DCN', 'SOX10', 'PRPH']
```
